# Bipolar and schizophrenia risk gene *AKAP11* encodes an autophagy receptor coupling the regulation of PKA kinase network homeostasis to synaptic transmission

**DOI:** 10.1101/2024.12.30.630813

**Authors:** You-Kyung Lee, Cong Xiao, Xiaoting Zhou, Le Wang, Meghan G. McReynolds, Xian Han, Zhiping Wu, Eric Purisic, Henry Kim, Xianting Li, Zhiping Pang, Jinye Dai, Junmin Peng, Nan Yang, Zhenyu Yue

## Abstract

Human genomic studies have identified protein-truncating variants in *AKAP11* associated with both bipolar disorder (BD) and schizophrenia (SCZ), implicating a shared disease mechanism driven by loss-of-function. AKAP11, a protein kinase A (PKA) adaptor, plays a key role in degrading the PKA-RI complex through selective autophagy. However, the neuronal functions of AKAP11 and the impact of its loss-of-function remains largely uncharacterized. Through multi-omics approaches, cell biology, and electrophysiology analysis in mouse models and human induced neurons, we delineated a central role of AKAP11 in coupling PKA kinase network regulation to synaptic transmission. Loss of AKAP11 distorted compartment-specific PKA and GSK3α/β activities and impaired cellular functions that significantly overlap with pathways associated with BD and SCZ. Moreover, we identified interactions between AKAP11, the PKA-RI adaptor SPHKAP, and the ER-resident autophagy-related proteins VAPA/B, which co-adapt and mediate PKA-RI complex degradation in neurons. Notably, AKAP11 deficiency impaired neurotransmission, providing key insights into the mechanism underlying *AKAP11*-associated psychiatric diseases.

## Introduction

A meta-analysis combining exome sequencing uncovered a link of rare protein-truncating variants (PTVs) to increased risk of bipolar disorder (BD) and identified *AKAP11* as a definitive risk gene for BD shared with schizophrenia (SCZ)^1^. Two independent meta-analyses of whole exome sequencing also revealed that *AKAP11*-coding or truncating variants enhanced the risk of SCZ across diverse human populations^2,3^. Thus, the human genetic evidence demonstrates *AKAP11* as a common risk gene for both BD and SCZ and implicates a potential disease mechanism involving the loss-of-function in the AKAP11. However, the biology and pathophysiology of AKAP11 in the central nervous system (CNS) remains poorly understood.

AKAP11 (A-kinase Anchoring Protein 11), known also as AKAP220, is a member of the protein family that anchors Protein Kinase A (PKA) complex to specific subcellular locations, modulating PKA and GSK-3β activity^4–8^. We previously reported a cellular function of AKAP11 as an autophagy receptor, which mediates the selective degradation of the regulatory RI subunits (RIα and RIβ) of PKA, while sparing RII subunits, through binding autophagy protein LC3 and controls PKA activity^9^. We recently documented the conserved autophagy function of AKAP11 in human neurons and demonstrated a critical role of AKAP11 in maintaining the homeostasis of PKA-RI protein complex in the soma and neurites through autophagy degradation^10^. Interestingly, *Akap11* knock-out mice exhibit behavioral abnormalities in electroencephalogram (EEG) recording in common with symptoms of SCZ^11^, supporting potential synaptic dysfunction^12^. Given the notion that neuronal autophagy regulates synaptic function^13–19^, it raises a question for how AKAP11-mediated autophagy is linked to the regulation of synaptic activity. The knowledge of the mechanism whereby AKAP11 controls PKA-RI complex homeostasis and synaptic functions in neurons is expected to provide insight into the pathogenesis of BD and SCZ.

Here we reported an interdisciplinary study of AKAP11 neuronal function and the consequence of the loss-of-function in AKAP11 in both genetic mouse models and human induced neurons (iNeurons) by integrating multi-omics, cell biology, and electrophysiology approaches. We observed an extensive distortion of PKA kinase network and altered PKA and GSK3α/β kinase activities in neurons from *AKAP11*-deficient neurons. Furthermore, loss of AKAP11 results in a compartment-specific change of PKA activity in neurons. Our study demonstrated the role of AKAP11 in coupling PKA kinase network regulation to synaptic transmission. Finally, our data identified a novel autophagy function of AKAP11 in neurons through interacting with the PKA adaptor SPHKAP and the ER-resident autophagy related proteins VAPA/B. Our current report offers a comprehensive view of *AKAP11*-regulated neuronal pathways and insight of the pathogenic mechanisms underlying the psychiatric diseases.

### Integrated proteomics identified profound changes of PKA kinase network and synaptic proteins in *AKAP11*-deficient neurons

We previously reported that *AKAP11* knockdown (KD) leads to an increase of protein level in the regulatory subunits RIα and RIβ of PKA in iNeurons^10^, confirming its role as autophagy receptor. Here we sought to determine AKAP11-targeted proteins and cellular pathways by performing a systematic search in neurons through integrated proteomics profiling, as we described in the study of autophagy cargo identification^10^(Fig. 1a). For this purpose, we established *Akap11*-conditional knock-out (cKO) (Cre-LoxP) and whole-body knock-out (wKO) mice (Extended Data Fig. 1a). Together with human *AKAP11*-KD iNeuron, we conducted quantitative mass spectrometry-based proteomics (Tandem Mass Tag (TMT) and Data-independent Acquisition (DIA))^20–22^ and analyzed the changes of proteins and cellular pathways in neurons from mutant mouse brains and iNeurons compared to control neurons. In total, 7427 proteins (54 up-regulated, 124 down-regulated), 5619 proteins (86 up-regulated, 29 down-regulated), and 11392 proteins (175 up-regulated, 13 down-regulated) were detected in *Akap11*-cKO, iNeuron-KD, and wKO datasets, respectively (Fig. 1b,c, Extended Data Fig. 1b and Supplementary Table 1-3).

**Fig. 1:**
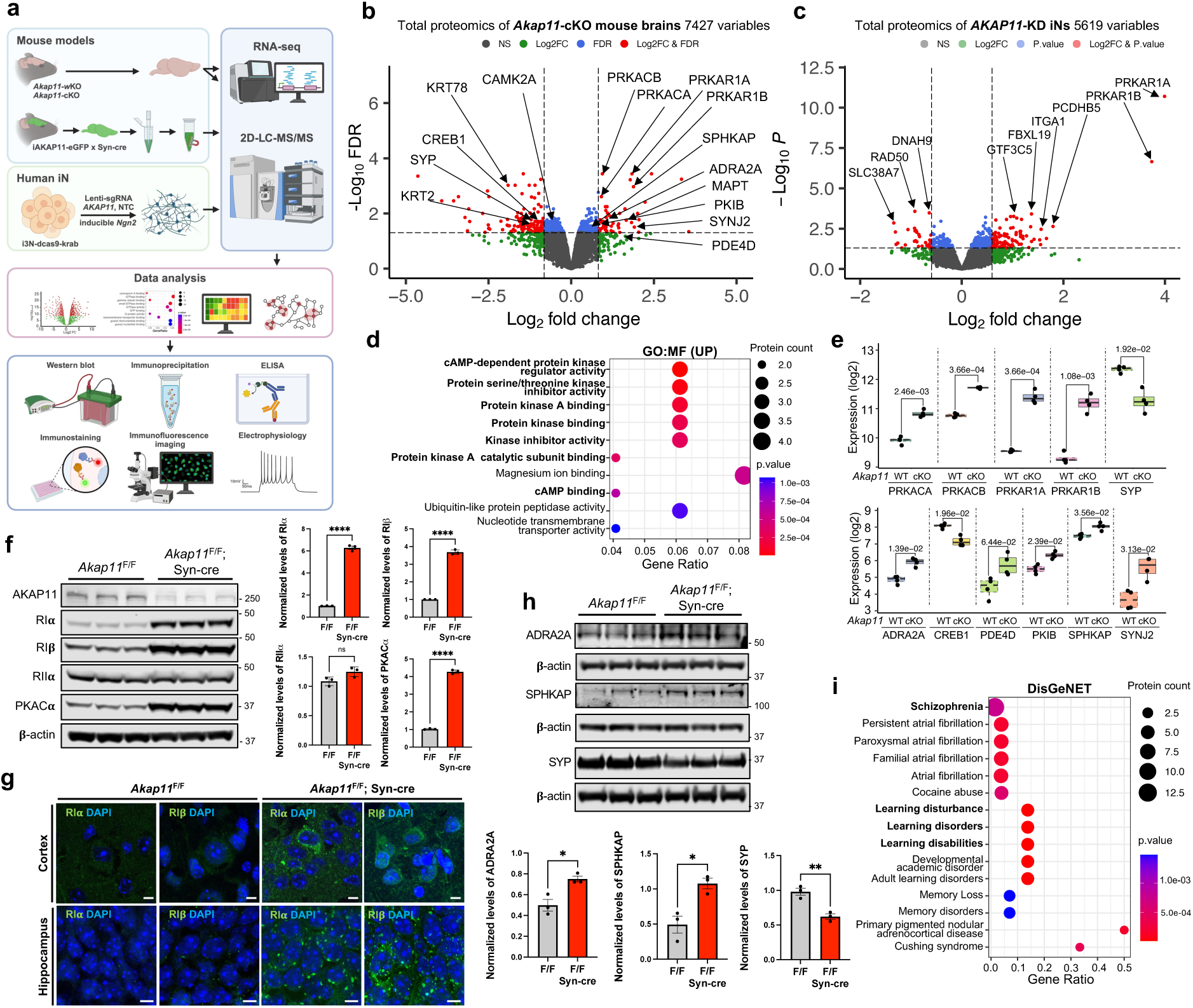
Integrated proteomic analysis of brain lysate from *Akap11*-cKO mouse brains and *AKAP11*-KD iNeurons. **a**, An overview of the study workflow, which integrates multiple-omics, bioinformatics, cell imaging, electrophysiology, and protein biochemistry, and investigates multiple genetic mouse models of AKAP11 and human iNeurons. Created with BioRender.com. **b**, Volcano plot of the differentially expressed proteins (DEPs) detected by mass-spectrometry in *Akap11*-cKO mouse brains. A positive score indicates enrichment, a negative score indicates depletion. The y axis represents statistical confidence (adjusted *P*-value (FDR)) for each x axis point. Enriched proteins, defined by FDR < 0.05 and |Log2FC| > 2SD, are colored in red. **c**, Volcano plot of the differentially expressed proteins (DEPs) detected by mass-spectrometry in *AKAP11*-KD human iNeurons. A positive score indicates enrichment, a negative score indicates depletion. The y axis represents statistical confidence (*P*-value) for each x axis point. Enriched proteins, defined by *P*-value < 0.05 and |Log2FC| > 2SD and followed by FDR estimation through permutation (FDR<0.05), are colored in red. **d**, Gene Ontology (GO) annotations of the DEPs in **b**, displaying the top 10 enriched pathways with *P*-value < 0.05. **e**, The enrichment levels for the proteins as shown in **b** with log2 transformed DIA intensities across *Akap11^F^*^/F^ and *Akap11*^F/F^; Syn-cre mice. **f**, Immunoblotting using antibodies against AKAP11 and PKA subunits and quantification of the blot results. Data are presented as mean ± SEM (n=3 per genotype). Statistical analysis was performed using an unpaired *t*-test. **g**, Immunofluorescence (IF) imaging of the cortex and hippocampus from *Akap11^F^*^/F^ and *Akap11*^F/F^; Syn-cre mice, stained with anti-RIα, anti-RIβ and DAPI. Scale bars, 5μm. **h**, Immunoblotting using antibodies against ADRA2A, SPHKAP, and SYP and quantification of the blot results. Data are presented as mean ± SEM (n=3 per genotype). Statistical analysis was performed using an unpaired *t*-test. **i**, DisGeNet annotations of the DEPs through ToppGene in **b** with *P*-value < 0.05.

We next annotated the differential expression of proteins (DEPs) in the *AKAP11*-cKO neurons. For the upregulated DEPs, functional annotation with clusterProfiler^23^ identified cAMP-dependent PKA binding related pathways as the dominant GO terms, including protein kinase regulator activity and PKA catalytic subunit binding under the Molecular Function (MF) category. In Cellular Component (CC) category, examples of significantly enriched GO terms include negative regulation of phosphorus metabolic process and negative regulation of kinase activity (Fig. 1d and Extended Data Fig. 1c). Indeed, we observed a significant enrichment of PKA-RIα (log2FoldChange (FC) 1.84) and RIβ (FC 1.88) levels as well as PKA-Cα (FC 0.91) and Cβ (FC 0.92) levels in *Akap11*-cKO dataset, and similar patterns in other two datasets (Fig. 1b,c,e and Extended Data Fig. 1b,d). Furthermore, multiple negative regulators of PKA activity are among the significantly upregulated DEPs in *Akap11*-cKO mouse brains, such as ADRA2A (FC 1.06), PDE4B (FC 1.38), PDE3A (FC 0.80), and PKIB (FC 0.83) (Fig. 1b,e and Extended Data Fig. 1b). ADRA2A inhibits adenyl cyclase (AC), leading to a decrease in protein kinase A (PKA) activity^24–26^. PDE3A and PDE4B inhibit PKA activity by breaking down cAMP, and PDE4B is a major binding partner of DISC1 and a risk gene for SCZ and BD^27–29^. PKIB belongs to cAMP-dependent protein kinase inhibitor family, and it regulates PKA pathway by interacting with the PKA-C subunit^30^. Additionally, we observed an enrichment of SPHKAP (FC 0.57, SD = 0.41), which is a PKA-RI subtype-specific anchoring protein^31^ (Fig. 1e). The increase of PKA-RIα/β, PKA-C (with a lesser degree than PKA-RI), ADRA2A, and SPHKAP proteins in *Akap11*-cKO brains were validated through immunoblot. The RIα/β proteins were accumulated in large discrete puncta in *Akap11*-deficient neurons as shown through immunofluorescence (IF) imaging (Fig. 1f,g,h).

In the downregulated DEPs, the annotation identified neural or synapse-related functions as the primarily affected terms, including synaptic vesicle (SV) membrane, exocytic vesicle membrane, SV, and myelin sheath (CC). Significant GO terms also include regulation of neurotransmitter transport and SV endocytosis in Biological Process (BP). Additionally, G protein activity, GTPase regulator activity, and transmembrane transporter binding are the main GO terms (MF) (Extended Data Fig. 1e). We noticed the downregulation of CREB1 (FC −0.92), a downstream target of PKA signaling pathway^32^, and synaptophysin (FC −1.09), a synaptic vesicle (SV)-associated protein important for synaptic trafficking and neurotransmitter release^33^. Additionally, the downregulated expression of calcium/calmodulin-dependent protein kinases, CAMKIIA (FC −0.51) indicates reduced neuronal calcium signaling, which plays a pivotal role in synaptic plasticity and memory formation^34^ (Fig. 1b).

To verify the changes of synaptic proteins identified in the proteomics analysis, we performed immunoblotting in *Akap11*-cKO brain lysates. We observed that the levels of synaptophysin were decreased in mutant brains (Fig. 1h). Using SynGO^35^ to screen the DEPs in *Akap11*-cKO mouse and *AKAP11*-KD iNeuron, we found an enrichment in both pre- and postsynaptic compartments, sharing multiple “sub-synaptic” terms, including regulation of synapse assembly, SV exocytosis and endocytosis, postsynaptic membrane neurotransmitter receptor levels, and chemical synaptic transmission (Extended Data Fig. 2a). The above evidence suggests that *AKAP11*-deficiency affects synaptic homeostasis and perhaps synaptic transmission.

We next performed transcriptome analysis of *Akap11*-cKO mouse brains. The number of differential expression genes (DEGs) in mutant mice is significantly less than that of DEPs (Extended Data Fig. 2b). Even with diminishing *Akap11* transcripts, *Akap11*-cKO brains showed little change of mRNA levels coding for PKA complex or its regulators as described above, supporting the regulatory role of AKAP11 in regulating PKA complex proteins post-transcriptionally. Meanwhile, Gene Set Enrichment Analysis (GSEA)^36,37^ indicates the down-regulated DEGs associated with synapse, including pre- and postsynaptic membrane, dendrite development, axonogenesis, actin and filament binding, etc. (Extended Data Fig. 2c).

Furthermore, through a comprehensive platform integrating information on human disease-associated genes and variants (DisGeNET)^38^, we found remarkably SCZ as the top ranked disease based on the protein count (Fig. 1i), suggesting a convergence of AKAP11-regulated cellular functions and potential SCZ-inflicted disease pathways.

### Identification of PKA subunits as central hubs in network modules for synapse formation regulated by AKAP11

To gain an insight of the relationship among cellular pathways/functions affected by the loss-of −function in *AKAP11* as autophagy receptor, we employed the modified Weighted Gene Co-expression Network Analysis (WGCNA)^39,40^ to dissect proteomics data from *Akap11*-cKO mouse brains and human *AKAP11*-KD iNeurons. WGCNA construction identified 13 and 18 modules in mutant mouse brains and human iNeurons, respectively (Fig. 2a,e).

**Fig. 2:**
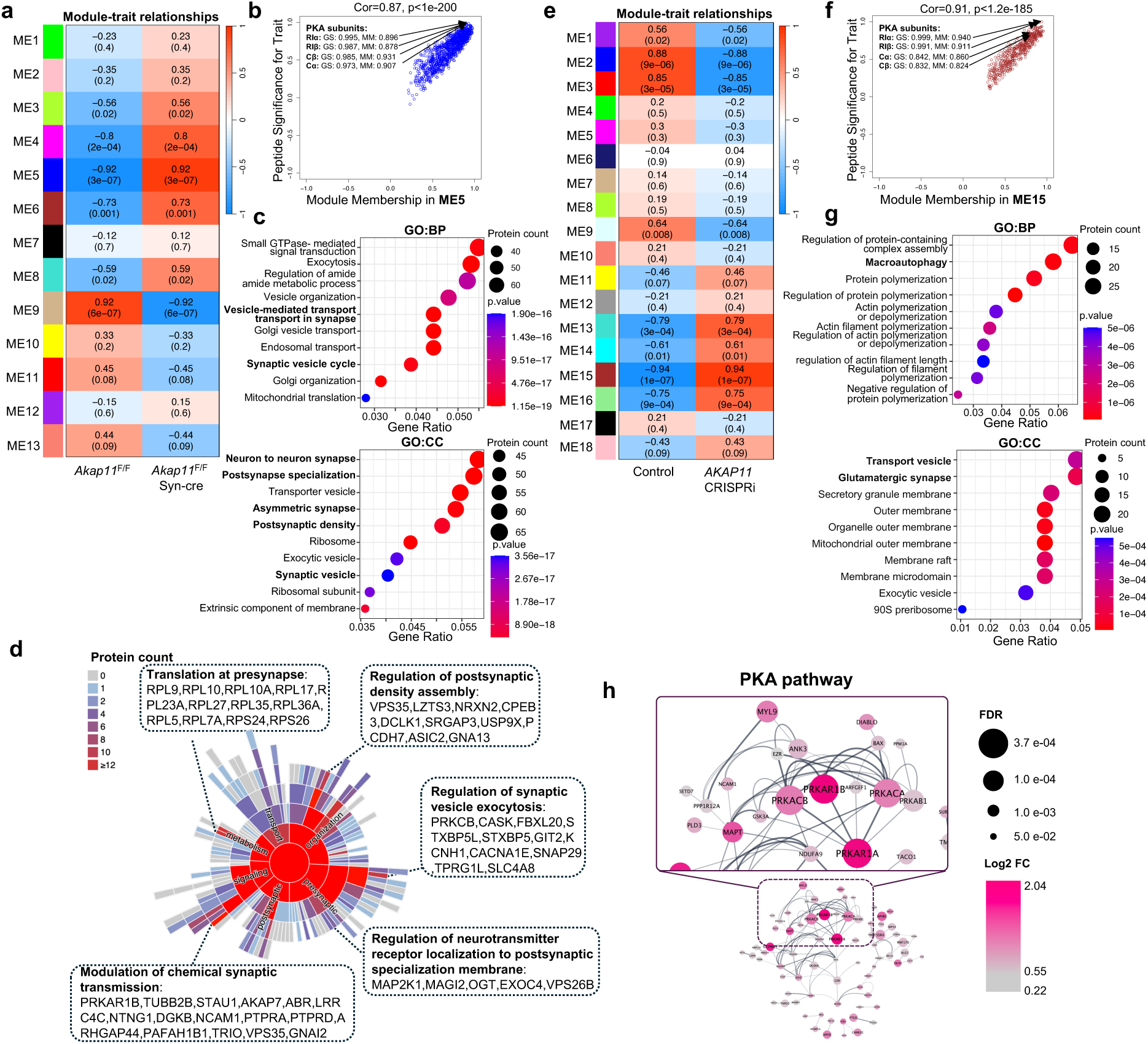
Identification of co-regulation of cellular pathways upon *AKAP11*-deficiency through WGCNA. **a**,**e**, Heatmap of the correlation between the module eigengenes and traits in *Akap11*-cKO mouse brains (16 samples) (**a**) or in *Akap11*-wKO iNeurons (16 samples) (**e**). Red represents positive correlation, and blue represents negative correlation to the trait. The *Pearson* correlation and *P*-value are presented in each module. **b**,**f**, Scatter plot of the module membership (x axis) and peptide significance (y axis) in ME5 from *Akap11*-cKO mouse brains (**b**) or ME15 from *AKAP11*-KD iNeurons (**f**). **c**,**g** Gene Ontology (GO) annotations of the proteins in ME5 from *Akap11*-cKO mouse brains (**c**) or ME15 from *AKAP11*-KD iNeurons (**g**), displaying the top 10 enriched pathways with *P*-value < 0.05. **d**, Sunburst plots depicting SynGO Biological Process pathways with a *P*-value < 0.05. For selected ontologies, representative proteins from ME5 within the ontology are displayed. **h**, PPI network analysis of proteins in ME5 in **a** (only FDR < 0.05). The color gradience represents from low (grey) to high (red) fold enrichment. Dot size represents statistical significance.

In *Akap11*-cKO brain dataset, module ME5 (1146 proteins) and ME4 (106 proteins) (Supplementary Table 4), are top ranked and positively correlated with the trait (mutant) with a *P*-value of 3e-07 and 2e-04, respectively. We found PKA-RIα/β and PKA-Cα/β are the hub proteins in the ME5 based on their highest scores in Peptide Significance (PS) and Module Membership (MM) (Fig. 2b). Interestingly, the GO term analysis reveals that ME5 is highly enriched in endosomal transport, postsynapse organization, synaptic vesicle and neuron to neuron synapse (Fig. 2c). Detailed analysis of “subsynaptic” processes of the ME5 showed the translation at presynapse, regulation of postsynaptic density assembly, SV exocytosis, chemical synaptic transmission, and neurotransmitter receptor localization of postsynaptic specialization membrane (Fig. 2d). In ME4, the proteins GOLPH3, PGM3, SSB, and CHMP2B are hub proteins and involved in protein metabolic process (Extended Data Fig. 3a). GO term analysis shows that ME4 is enriched in protein catabolic process, cytoplasmic translation, and proteasome complex (Extended Data Fig. 3b).

In *AKAP11*-KD iNeuron dataset, the ME15 (482 proteins) and ME13 (344 proteins) are the top two positively correlated modules (Supplementary Table 5). Similarly, PKA-RIα/β subunits were identified as hub proteins followed by PKA-Cα/β in the ME15 (significant PS and MM), while the ME15 showed the enrichment in macroautophagy, glutamatergic synapse, transport vesicle, and protein polymerization or depolymerization (Fig. 2f,g). In ME13, NUP35, NMT2, MTM1, and BCL2 are hub proteins, which are involved in endomembrane system, while GO term analysis reveals the association of ME13 with endosomal transport, early endosome, and lysosomal membrane (Extended Data Fig. 3c,d). Therefore, the data from both mouse brains and human iNeuron show a remarkable consensus where PKA subunits are the hub proteins in the synapse-related modules. The results demonstrate a functional accordance of PKA signaling pathways with synaptic function, which are co-regulated by AKAP11. Moreover, the data indicated a functional correlation of PKA signaling, autophagy, and vesicle trafficking pathway through AKAP11, in agreement with our early observation^9^.

We next employed STRING^41^ to analyze protein-protein interaction (PPI) network of the PKA-centered protein module (ME5) (only FDR < 0.05 proteins). While PKA-RIα/β as the most significant DEPs (increase) in *Akap11*-cKO brains and as the main hub proteins (upregulated) in the ME5 (Fig. 1b), they are found as the central nodes in the PPI network, which interact with MAPT, GSK3A, ARFGEF1, ANK3, EZR, NDUFA9, and PRKAB1 (Fig. 2h). Indeed, those interacting proteins are the known PKA targets or mediators of PKA signaling pathways in various cellular functions or neural activity^6,31,42–50^. The evidence indicated a significant alteration of PKA signaling network, including PKA holoenzyme subunits, upstream regulators, and downstream targets, in *AKAP11*-deficient neurons.

### Disruption of neuronal AKAP11 altered protein phosphorylation landscape and compartment-specific PKA activities

The increase of protein levels in PKA regulatory subunits RIα/β as well as in multiple PKA activity inhibitors (Fig. 1e,f,h and Extended Data Fig. 1d) led to a hypothesis that PKA activity is disrupted in *AKAP11*-deficient neurons. To test the hypothesis, we analyzed phosphoproteomics data obtained from the brains of *Akap11*-cKO mice and determined changes of potential PKA phosphorylation sites. We identified 289 differentially abundant phosphoprotein sites (DAPPSs) between the *Akap11*-cKO and control brain lysates (91 upregulated and 198 downregulated) (Fig. 3a and Supplementary Table 6). Strikingly, given the finding of genetic risk of *AKAP11* variants in SCZ and BD, the enrichment analysis of the DAPPSs corresponding proteins through the DisGeNET^38^ database reveals an enrichment of diseases such as SCZ, BD, intellectual disability, neurodevelopmental disorders, and unipolar depression in *Akap11-*cKO brains (Fig. 3b).

**Fig. 3:**
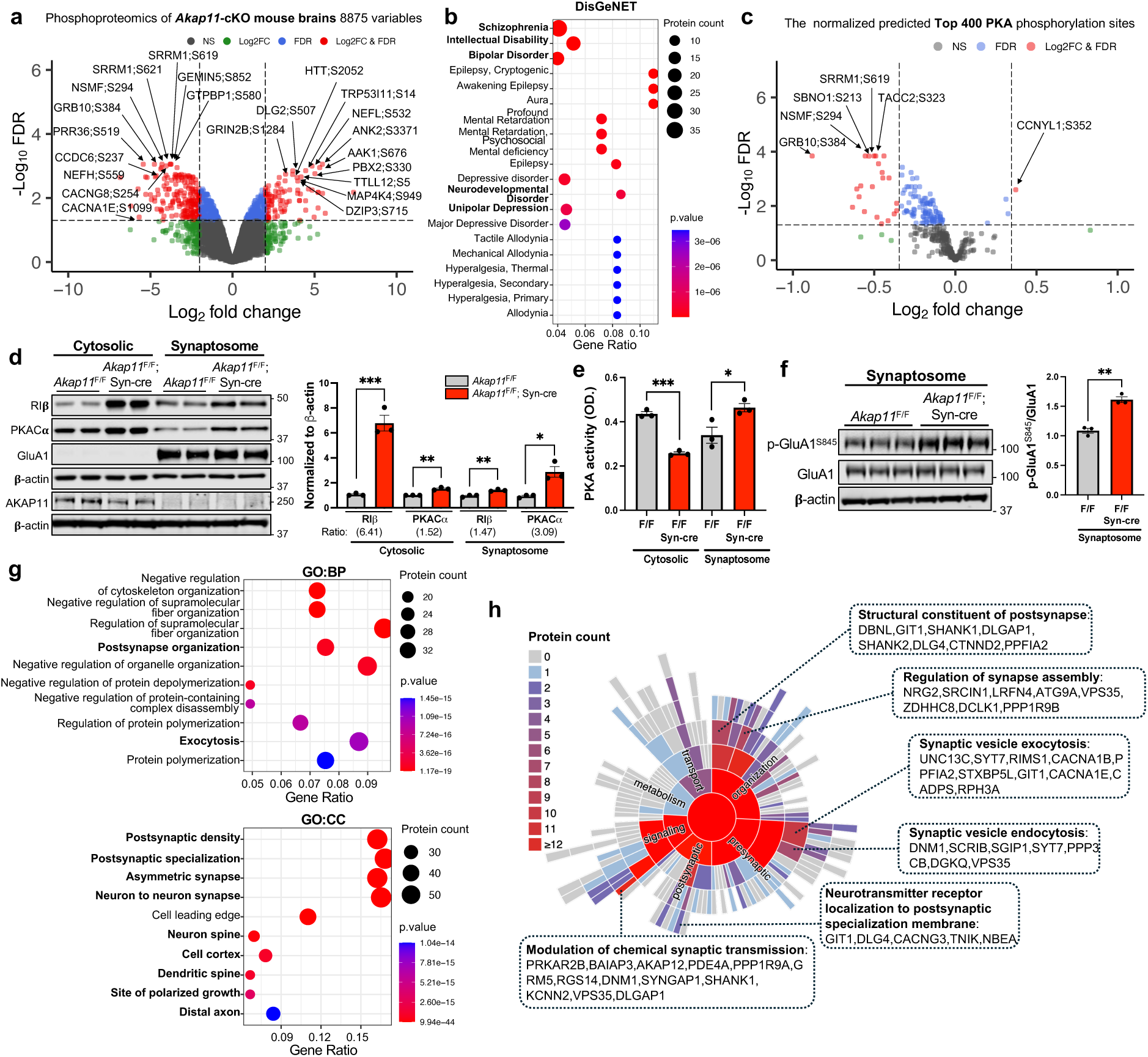
Phosphoproteomics and PKA activity analysis of *Akap11-*cKO brain lysates. **a**, Volcano plot of the differentially abundant phosphoprotein sites (DAPPSs) detected by mass-spectrometry in *Akap11*-cKO mouse brains and *AKAP11*-KD human iNeurons. A positive score indicates enrichment, a negative score indicates depletion. The y axis represents statistical confidence (adjusted *P*-value (FDR)) for each x axis point. Enriched proteins, defined by FDR < 0.05 and |Log2FC| > 2SD, are colored in red. **b**, DisGeNet annotations of the DEPPSs through ToppGene in **a** with *P*-value < 0.05. **c**, Volcano plot of the top 400 predicted PKA phosphorylation sites (normalized by the total protein level) detected by mass-spectrometry in *Akap11*-cKO mouse brains. A positive score indicates enrichment, a negative score indicates depletion. The y axis represents statistical confidence (adjusted *P*-value (FDR)) for each x axis point. Enriched proteins, defined by FDR < 0.05 and |Log2FC| > 2SD, are colored in red. **d,** Immunoblotting using antibodies against RIβ, PKA-Cα, GluA1, and AKAP11 in cytosolic and synaptosome fractions and quantification of the results. Data are presented as mean ± SEM (n=3 per genotype). Statistical analysis was performed using an unpaired *t*-test. **e**, Bar graphs of the ELISA results measuring PKA activity in the cytosol and synaptic fractions from *Akap11^F^*^/F^ and *Akap11*^F/F^; Syn-cre mice. Data are presented as mean ± SEM. Statistical analysis using an unpaired *t*-test. (n=3 per genotype). **f,** Immunoblotting using antibodies against p-GluA1^S845^ and GluA1 in synaptosome and quantification of the results. Data are presented as mean ± SEM (n=3 per genotype). Statistical analysis was performed using an unpaired *t*-test. **g**, Gene Ontology (GO) annotations of the predicted PKA substrates, displaying the top 10 enriched pathways with *P*-value < 0.05. **h**, Sunburst plots depicting SynGO Biological Process pathways with a *P*-value < 0.05. For selected ontologies, representative proteins from predicted PKA substrates within the ontology are displayed.

We next employed a comprehensive pipeline PhosR^51^ to delineate the kinase and substrate relationship. The PhosR utilizes the PhosphoSitePlus^52^ as a reference database and a set of tools and methodologies implemented in R to allow the comprehensive analysis of the phosphoproteomic data^53^. The predicted top three phosphorylation sites of the corresponding kinases are shown in a heatmap (Extended Data Fig. 4a). Additionally, we analyzed the top 400 predicted PKA phosphosites and found 39 significantly downregulated DAPPSs and 41 phosphorylation sites with a trend of decrease (Extended Data Fig. 4b). After normalizing against the levels of corresponding proteins from proteomics results (Supplementary Table 1), we identified 22 downregulated DAPPSs (Fig. 3c) associated potentially with PKA activity in *Akap11*-cKO neurons.

In a parallel analysis, we mapped the phosphoproteomic data against the PKA consensus substrates from PhosphoSitePlus^52^ and identified 30 phosphosites (of which 20 were found as predicted PKA phosphosites from the PhosR analysis). Among them, 6 show a trend of increase while 9 display a trend of reduction (FDR <0.2, 1*SD) (Extended Data Fig. 4c).

To validate PKA activity changes, we first assessed the protein levels of PKA subunits in cytosol and synaptosome fractions and found an increase of PKA-C and PKA-RI in both fractions from *Akap11*-cKO brain lysates by immunoblot. We noticed that the increase of RΙβ was much greater in cytosol (6.41-fold) than in synaptosome (1.47-fold), while PKA-C changed in the opposite direction within cytosolic (1.52-fold) and synaptosome (3.09-fold) (Fig. 3d). We then performed an enzyme-linked immunosorbent assay (ELISA) to analyze PKA activity^54,55^. The assay sensitivity was verified using forskolin (FSK, a PKA activator) and H89 (a PKA inhibitor) (Extended Data Fig. 4d). Remarkably, the ELISA results showed a reduction PKA activity in the cytosol but an increase in the synaptosome (Fig. 3e). The increase of PKA activity in the synaptosomes were supported by increased phosphorylation of the PKA substrate p-GluA1^S845^ and of PKA-Cα^T197^ (Fig. 3f and Extended Data Fig.4e). The reduced PKA activity in the cytosol occurred in different brain regions, such as the cerebral cortex and hippocampus, in *Akap11*-wKO mice (Extended Data Fig. 4f). These findings indicate that the loss of AKAP11 induces compartment-specific alterations of PKA activities. It is conceivable that the greater elevation of regulatory subunit of PKA-RI levels causes repression of PKA activity in the cytosol, while the increase of PKA-C in the synaptosome fraction, which escape from binding to a moderate enhancement of PKA-RI, could result in the increase of PKA activity in synapse.

Moreover, we performed the GO term analysis of the 399 predicted PKA substrates based on PhosR (Supplementary Table 7), the top enriched pathways were postsynaptic density and specialization, asymmetric synapse, and neuron spine axon (CC), post-synapse organization and exocytosis (BP), and actin binding (MF) (Fig. 3g and Extended Data Fig. 4g). The SynGO^35^ analysis further showed the enrichment of the predicted PKA substrates in structural constituent of postsynapse, regulation of synapse assembly, SV exocytosis and endocytosis, and modulation of chemical synaptic transmission (Fig. 3h). These evidences corroborate the results from WGCNA showing the accordance of PKA network with synaptic activity (Fig. 2c,d,h).

### Loss-of-function in AKAP11 impaired GSK3α/β kinase activity in neurons

The PhosR further predicted a reduction in the kinase activities of GSK3β, PKC isoforms (α, γ, 8, ι, θ), and CaMKIIα alongside PKA, while AKT1 activity exhibits an increase (Fig. 4a).

**Fig. 4:**
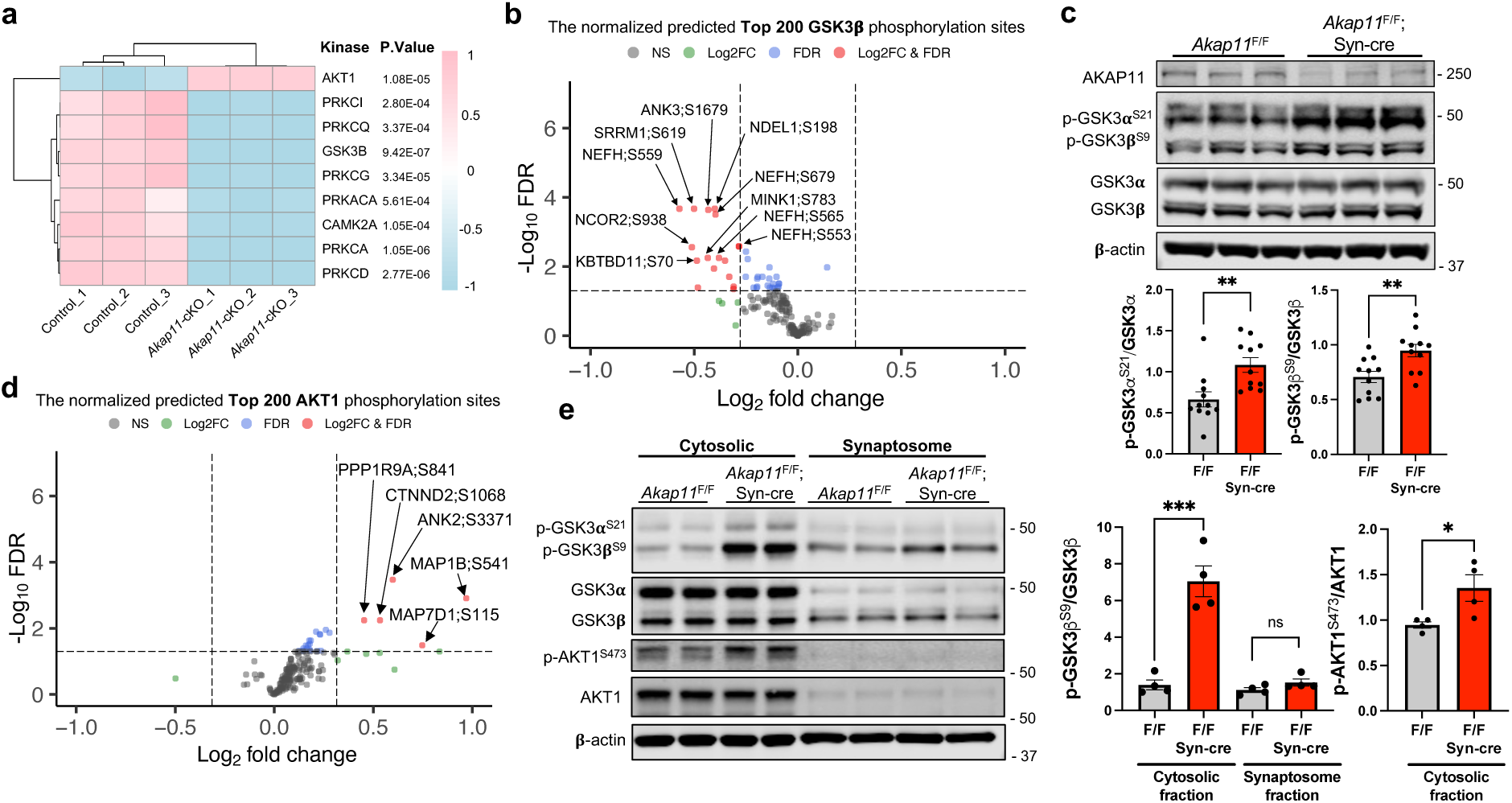
GSK3α/β kinase activity analysis in *Akap11*-cKO brain lysates. **a**, The predicted kinase activity scores for representative kinases. The scores were generated by PhosR based on the phosphorylation levels of predicted substrates. Statistical significance was determined using *t*-test. **b**, Volcano plot of the top 200 predicted GSK3β phosphorylation sites (normalized by the total protein level) detected by mass-spectrometry in *Akap11*-cKO mouse brains. A positive score indicates enrichment, a negative score indicates depletion. The y axis represents statistical confidence (adjusted *P*-value (FDR)) for each x axis point. Enriched proteins, defined by FDR < 0.05 and |Log2FC| > 2SD, are colored in red. **c**, Immunoblotting using antibodies against AKAP11, GSK3α/β and p-GSK3α/β (S21/S9) and quantification of the results. Data are presented as mean ± SEM (n=11 per genotype). Statistical analysis was performed using an unpaired *t*-test. **d**, Volcano plot of the top 200 predicted AKT1 phosphorylation sites (normalized by the total protein level) detected by mass-spectrometry in *Akap11*-cKO mouse brains. A positive score indicates enrichment, a negative score indicates depletion. The y axis represents statistical confidence (adjusted *P*-value (FDR)) for each x axis point. Enriched proteins, defined by FDR < 0.05 and |Log2FC| > 2SD, are colored in red. **e,** Immunoblotting using antibodies against p-GSK3α/β (S21/S9), GSK3α/β, p-Akt1^S473^, and Akt1 in cytosolic and synaptosome fractions and quantification of the results. Data are presented as mean ± SEM (n=4 per genotype). Statistical analysis was performed using an unpaired *t*-test.

Because GSK3α/β are known AKAP11-binding proteins^5^, we then analyzed GSK3α/β activities using PhosR and examined the specific phosphorylation sites of the kinase as a proxy through immunoblot. From the top 200 predicted GSK3β sites normalized by corresponding protein levels from proteomics results, we identified 17 DAPPSs, which were all downregulated (Fig. 4b), suggesting a decrease of GSK3β activity in *Akap11*-cKO neurons. Immunoblot with anti-phospho-GSK3α/β antibodies (Ser21 in GSK3α and Ser9 in GSK3β) showed enhanced phosphorylation levels of GSK3α/β in both cytosolic and synaptosome fractions in *Akap11*-cKO mouse brain (Fig. 4c,e) as well as *Akap11*-whole body KO mouse brains (Extended Data Fig. 5a). Our findings revealed that both AKAP11 and GSK3α/β were predominantly distributed in the cytosol relative to synaptosome fractions (Fig. 3d,4e), which aligns with the report that AKAP11directly binds GSK3α/β^5^. These above results demonstrate a reduction of GSK3α/β activities in *AKAP11* deficient neurons.

Additionally, we analyzed AKT1 kinase activity based on the normalized predicted top 200 AKT1 substrates. We identified 5 DAPPSs, all of which were upregulated. Among the rest of the predicted substrates, many exhibited a trend of increased phosphorylation levels (Fig. 4d). Immunoblot of the p-AKT1 Ser473 with a specific antibody, which detects phosphorylation of Ser 473 within the carboxyl-terminal hydrophobic motif that is required for Akt/PKB activation^56^, showed an increase in the cytosolic fraction in *Akap11*-cKO neurons (Fig. 4e). Note that p-AKT1 is nearly undetectable in the synaptosome fraction (Fig. 4e). The results corroborate an increase in AKT1 activity in the cytosol in *Akap11*-cKO neurons. Given that AKT1 can phosphorylate GSK3α (Ser 21) and GSK3β (Ser 9)^57^, our data are consistent with an idea that the loss of AKAP11 causes AKT1 activation, leading to the elevated phosphorylation of unbound GSK3α/β kinase and their inhibition.

Furthermore, our expanded analysis of the kinase network and substrate module pattern showed that *AKAP11*-deficiency causes a distortion of PKA, GSK3α/β, and AKT1 activities in neurons (Extended Data Fig. 5b,c). Taken together, our results reveal a fundamental role of AKAP11 in controlling PKA, GSK3a/b, and AKT1 activities in neurons.

### AKAP11-interactome analysis identifies new partners in PKA-RI adapting and autophagy function

To further elucidate AKAP11 function, we investigated the interactome of AKAP11 in CNS neurons. To this end, we generated transgenic mice with Cre-dependent expression of AKAP11-eGFP fusion protein. Crossing the transgenic with Synapsin-Cre mice enabled neuron-specific expression of AKAP11-eGFP fusion in mouse brain (Fig. 5a). Western blotting confirmed the expression of the AKAP11-eGFP fusion protein in the mice with AKAP11-eGFP allele carrying Synapsin-cre (Extended Data Fig. 5d). In addition, AKAP11-eGFP co-localizes with anti-AKAP11 antibody signals in the discrete puncta in neurons (Fig. 5b). Examination of the transgenic showed a wide distribution of the fluorescent AKAP11 fusion in the neurons across mouse brain. AKAP11-eGFP was expressed strongly in the cortex (particularly in layers 2 and 5/6), hippocampus (CA3 and DG), and the thalamus, and detected in the pons, and the cerebellum (e.g. Purkinje neuron) (Fig. 5c).

**Fig. 5:**
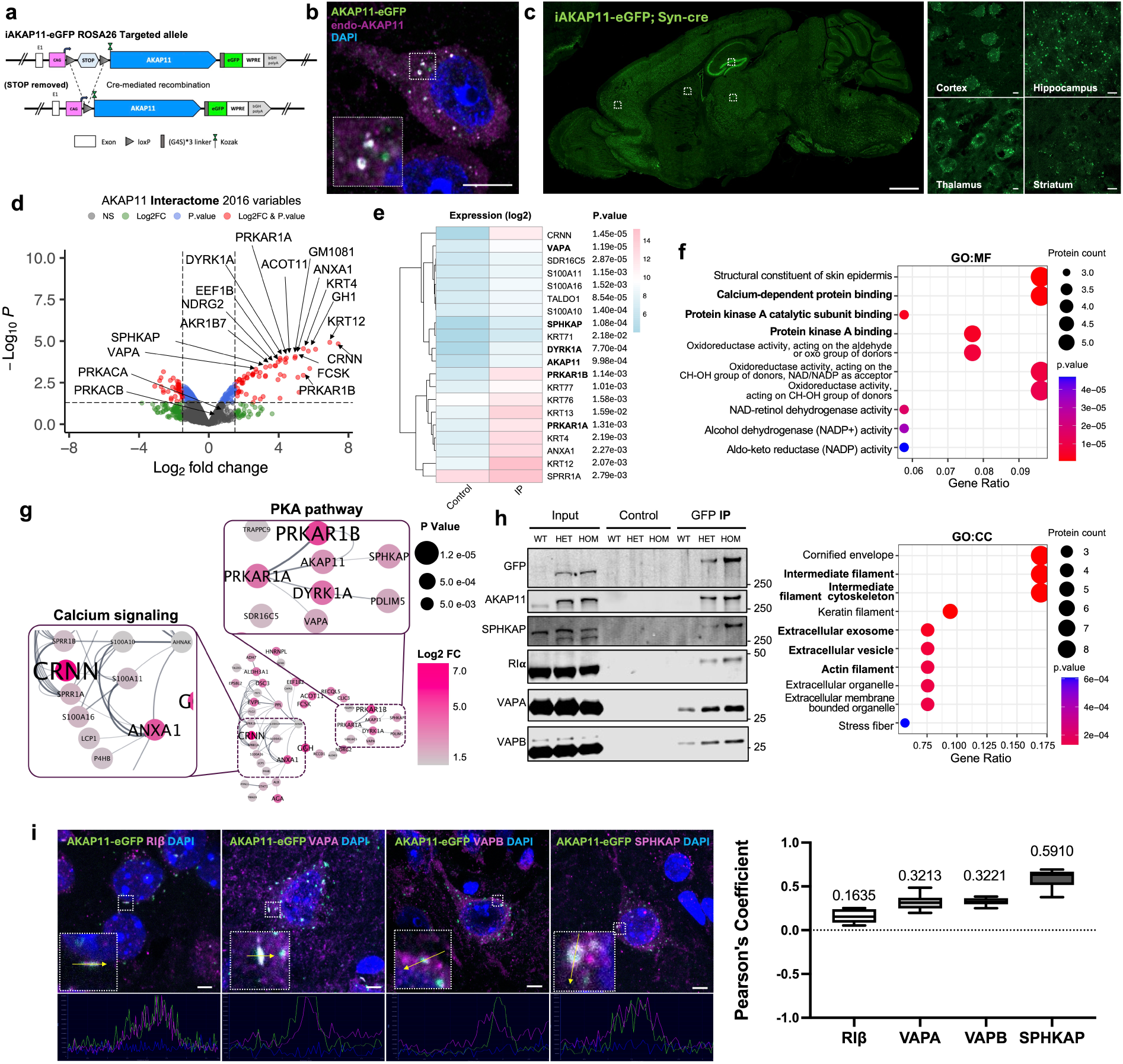
Characterization of AKAP11 interactome in mouse CNS neurons. **a**, Schematic illustration of the generation of AKAP11-eGFP; Syn-cre mice producing the fusion protein in CNS neurons. **b**, IF images of AKAP11-eGFP; Syn-cre brain slices stained with anti-GFP (green), anti-AKAP11 antibodies (magenta), and DAPI. Scale bar, 5μm. **c**, IF images of sagittal sections from AKAP11-eGFP; Syn-cre mouse brains, showing AKAP11 expression in various brain regions. Inset images provide magnified views of cortex, hippocampus, thalamus, and striatum. Scale bars, 1000μm (main images); 5μm (inset images). **d**, Volcano plot of the co-Immunoprecipitated (IP) proteins detected by mass-spectrometry in AKAP11-eGFP mouse brain lysates. A positive score indicates enrichment, a negative score indicates depletion. The y axis represents statistical confidence (*P*-value) for each x axis point. Enriched proteins, defined by *P*-value < 0.05 and |Log2FC| > 1.5SD and followed by FDR estimation through permutation (FDR<0.05), are colored in red. **e**, Enrichment levels of the specific proteins in **d** with log2 transformed DIA intensities across control and AKAP11-eGFP groups. **f**, Gene Ontology (GO) annotations of the proteins shown in **d**, displaying the top 10 enriched pathways with *P*-value < 0.05. **g**, PPI network analysis of proteins shown in **d**. The color gradience represents from low (grey) to high (red) fold enrichment. Dot size represents statistical significance. **h**, IP was performed using Chromotek GFP-trap® particle from heterozygous and homogenous of AKAP11-eGFP transgenic mice and non-transgenic mouse brains and immunoblotting with antibodies against GFP, AKAP11, SPHKAP, RIα, VAPA, and VAPB. **i,** Representative IF images of AKAP11-eGFP mouse brain slices stained with anti-RIβ, anti-VAPA, anti-VAPB, and anti-SPHKAP antibodies in the cortex, showing co-localization of these proteins. Scale bars, 5μm. Co-localization of AKAP11-eGFP with RIβ, VAPA, VAPB, and SPHKAP quantified using Pearson’s correlation coefficient calculated with the JACoP plugin. Data are presented as mean ± SEM (n >8 per group).

We then performed anti-GFP affinity purification, visualized by silver staining (Extended Data Fig. 5e), and conducted mass spectrometry analysis of co-purified proteins with AKAP11. The anti-GFP-bead uncovered over 2000 proteins, of which 59 were significantly enriched compared to the control. As anticipated, we observed significant enrichments of PKA RIα/β (FC 4.08/5.41) (Fig. 5d,e and Supplementary Table 8). The GO terms of the significantly enriched proteins highlight protein kinase A binding and calcium-dependent protein binding (MF), intermediate filament cytoskeleton, extracellular exosome and vesicle, and actin filament (CC), cytoskeleton organization and intermediate filament-based process (BP) (Fig. 5f and Extended Data Fig. 5f). The PPI analysis reveals two distinct clusters, including PKA pathway and calcium signaling, with AKAP11, PKA-RIα/β, SPHKAP, VAPA, and DYRK1A forming an interacting network within PKA cluster, and ANXA1 emerging as the central node in the calcium-dependent protein binding cluster (Fig. 5g). The identification of SPHKAP as a AKAP11-binding protein is intriguing, as SPHKAP protein levels were significantly increased in *AKAP11*-deficient neurons, and it exhibits a strong accordance with PKA subunits within the same protein module (Fig. 1b,2h). Moreover, VAP proteins are endoplasmic reticulum (ER)-resident proteins that directly interact with multiple autophagy-related (ATG) proteins. The interaction between ATG proteins and VAPA/B is a prerequisite for autophagosome formation^58,59^. Furthermore, the interaction between VAPA/B and SPHKAP was recently shown to result in the concentration of PKA-RI complex between stacked ER cisternae associated with ER-PM junctions, which is important for the reciprocal regulation of PKA and Ca2+ signaling and the coupling of excitation and transcription^60,61^. Through co-IP experiments we verified the interactions of AKAP11-eGFP with RIα/β, SPHKAP, and VAPA/B (Fig. 5h) in mouse brains with immunoblot analysis.

To understand the nature of the interactions between AKAP11, SPHKAP, and VAPA/B in neurons, we next examined their subcellular localization using the transgenic mice expressing AKAP11-eGFP (Extended Data Fig. 6a). Immunofluorescence (IF) imaging showed co-localization of AKAP11-eGFP with the endogenous RIα/β, SPHKAP, and VAPA/B in a fraction of discrete puncta in mouse CNS neurons (Fig. 5i and Extended Data Fig. 6b). Note the low score of the *Pearson*’s coefficient for RIβ, which could reflect the poor specificity of anti-RIβ antibody. Given VAPA/B as ER-residents^62,63^ and SPHKAP-VAPA/B complex known for adapting of PKA-RI at the ER-PM junction^61^, it raises a possibility that both AKAP11 and PKA-RI are recruited to the ER through interacting with SPHKAP and/or VAPA/B. We also observed a partial co-localization of AKAP11-eGFP with autophagy adaptors p62/SQSTM1 and LC3B in neurons, in agreement with its role in autophagy adapting (Extended Data Fig. 6b).

Together, the above findings showed the AKAP11 interactome associated with various cellular pathways. Importantly, the results from the analysis of PKA cluster offer additional support to the function of AKAP11 in regulating PKA protein homeostasis and autophagy process likely through interacting with SPHKAP and VAPA/B adaptor proteins, which are involved in adapting PKA-RI and autophagosome biogenesis^58,60,61^.

### Degradation of PKA adaptor SPHKAP depends on AKAP11-mediated autophagy

We found that the overall staining intensity of SPHKAP protein was markedly enhanced in the cortex and hippocampus of *Akap11*-cKO brains, compared to the control. For instance, a remarkable accumulation of SPHKAP protein was seen in the CA2, CA3, and DG of the hippocampus in *Akap11*-cKO brain (Fig. 6a and Extended Data Fig. 6c). High magnification of IF images showed that the SPHKAP puncta was significantly larger and more abundant in the neurons of *Akap11*-cKO brains than those in control mouse brains (Fig. 6a). Considering the increase of SPHKAP protein levels in *Akap11*-cKO brain (Fig. 1b), the observation suggests that the loss of AKAP11 disrupts the SPHKAP homeostatic levels in neurons.

**Fig. 6:**
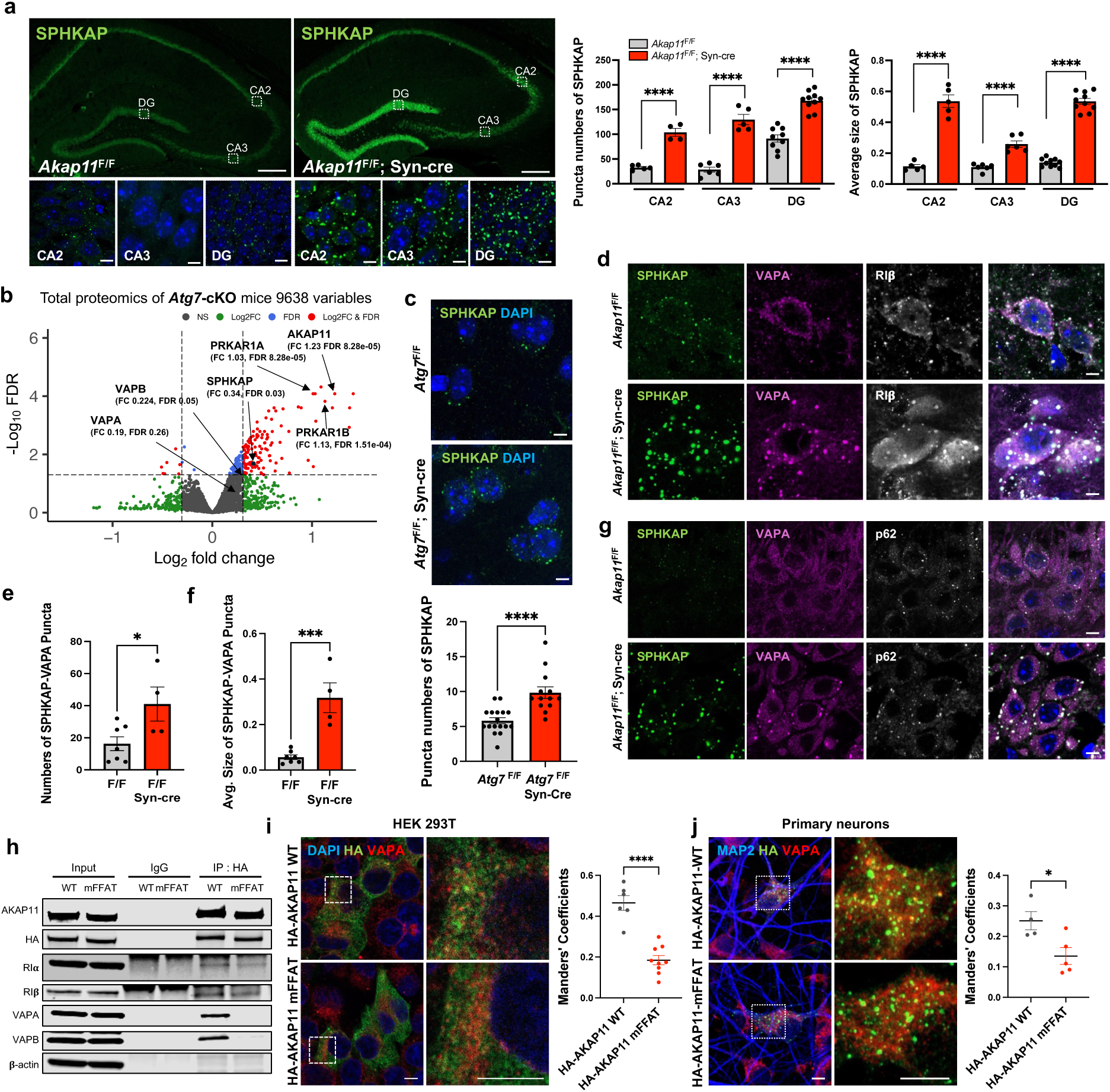
Characterization of SPHKAP and VAP proteins in neurons from *Akap11*-cKO and *Atg7*-cKO mouse brains. **a,** IF imaging of the hippocampal CA2, CA3, and DG of the brains slices from *Akap11^F^*^/F^ and *Akap11*^F/F^; Syn-cre mice, stained with anti-SPHKAP and DAPI. Inset images provide magnified views of cortex, hippocampus, thalamus, and striatum. Scale bars, 200μm (main images); 5μm (inset images). Bar graph as the quantification of SPHKAP puncta numbers and average size in DAPI-positive cells using Analyze Particles in ImageJ. Data are presented as mean ± SEM (n > 4 per genotype). Statistical significance (****p<0.0001) was performed using an unpaired *t*-test. **b**, Volcano plot of the differentially expressed proteins (DEPs) detected by mass-spectrometry in *Atg7^F^*^/F^ and *Atg7*^F/F^; Syn-cre mouse brains. A positive score indicates enrichment, a negative score indicates depletion. The y axis represents statistical confidence (adjusted *P*-value (FDR)) for each x axis point. Enriched proteins, defined by FDR < 0.05 and |Log2FC| > 2SD, are colored in red. **c**, IF imaging of the cortex from *Atg7^F^*^/F^ and *Atg7*^F/F^; Syn-cre mice, stained with anti-SPHKAP and DAPI. Scale bars, 5 μm. Bar graph as the quantification of SPHKAP puncta numbers per DAPI-positive cell using Analyze Particles in ImageJ. Data are presented as mean ± SEM (n > 13 per genotype). Statistical significance (****p<0.0001) was performed using an unpaired *t*-test. **d,e,f,** IF imaging of the cortex from *Akap11^F^*^/F^ and *Akap11*^F/F^; Syn-cre mice, stained with anti-SPHKAP, anti-VAPA, anti-RIβ, and DAPI. Scale bars, 5 μm. Bar graph as the quantification of SPHKAP-VAPA puncta numbers and average size in DAPI-positive cells using Analyze Particles in ImageJ. Data are presented as mean ± SEM (n >4 per genotype). Statistical significance (*p<0.05 and ***p<0.001) was performed using an unpaired *t*-test. **g**, IF imaging of the cortex from *Akap11^F^*^/F^ and *Akap11*^F/F^; Syn-cre mice, stained with anti-SPHKAP, anti-VAPA, anti-p62, and DAPI. Scale bars, 5 μm. **h,** Co-immunoprecipitation was performed using anti-HA antibody from HEK293T cells expressing HA-tagged AKAP11 WT or FFAT motif mutant (mFFAT), followed by immunoblotting with antibodies against HA, AKAP11, RIα, RIβ, VAPA, and VAPB. **i,** HEK293T cells expressing HA-AKAP11 WT or mFFAT were stained with antibodies against HA, VAPA, and DAPI. The bar graph shows quantification of HA–VAPA co-localized signal using Manders’ coefficient. Data are presented as mean ± SEM (n > 6 per group). Statistical significance (****p<0.0001) was performed using an unpaired *t*-test. Scale bars, 5 μm. **j,** Primary neurons expressing HA-AKAP11 WT or mFFAT were stained with antibodies against HA, VAPA, and MAP2. The bar graph shows quantification of HA–VAPA co-localized signal using Manders’ coefficient. Data are presented as mean ± SEM (n > 4 per group). Statistical significance (*p<0.05) was performed using an unpaired *t*-test. Scale bars, 5 μm.

Given the role of AKAP11 as autophagy receptor in PKA-RI degradation^9,10^, we next asked if SPHKAP, like PKA-RI, is selectively degraded through autophagy. Examination of our previous proteomics study of *Atg7*-cKO brains (deficient in autophagy)^10^ showed a significant enrichment of SPHKAP, RIα/β, and AKAP11 protein levels (but not VAP proteins) (Fig. 6b). Furthermore, SPHKAP protein was accumulated in *Atg7*-cKO neurons with increased number of puncta compared to the control (Fig. 6c). Thus, the evidence from both *Akap11*-cKO (Fig. 1b) and *Atg7*-cKO neurons suggests that AKAP11 mediates selective degradation of SPHKAP through autophagy.

To ask if AKAP11 regulates the function of SPHKAP and VAPA/B in adapting PKA-RI at ER, we assessed the subcellular localization of SPHKAP, VAPA/B, and PKA-RI in *Akap11*-cKO brains. Remarkably, IF staining showed that SPHKAP and VAPA were accumulated and colocalized in the large puncta, with a significant increase in both the SPHKAP-VAPA puncta number and size in the neurons of *Akap11*-cKO brains, compared to the control. RIβ exhibited extensive co-localization with SPHKAP-VAPA puncta (Fig. 6d,e,f). The results suggest that the loss of AKAP11 does not disrupt the interactions between SPHKAP, VAP proteins, and RIβ. Instead, lacking AKAP11 resulted in the expansion of the SPHKAP-VAPA-RIβ puncta structures, suggesting an impairment of the clearance. Furthermore, IF staining showed a partial colocalization of p62 signal with the SPHKAP-VAPA puncta, evidenced in the hippocampus from *Akap11*-cKO brain (Fig. 6g). Similar results of the enhanced co-localization for the three proteins were observed in primary neurons from *Akap11*-wKO mice (Extended Data Fig. 6d,e). Taken together, these data suggest that AKAP11 is not required for the function of SPHKAP/VAP proteins in adapting PKA-RI to ER; rather AKAP11 mediates the degradation of SPHKAP alongside PKA-RI through selective autophagy.

To evaluate whether AKAP11 regulates VAPA degradation through direct interaction, we generated a mutant AKAP11 lacking a functional FFAT motif, which is known to mediate binding to VAP proteins^60^. Immunoprecipitation (IP) showed that the FFAT-mutant AKAP11 failed to interact with VAPA and VAPB, while wild-type AKAP11 exhibited robust binding in transfected HEK293T cells (Fig. 6h). In addition, co-localization between AKAP11 and VAPA was markedly disrupted by the FFAT mutant, as shown by IF imaging in both HEK 293T cells and primary neurons (Fig. 6i,j and Extended Data Fig. 6f). These findings suggest that the FFAT motif is required for the AKAP11-VAPA interaction, potentially enabling the recruitment of the AKAP11 to the ER.

### Impaired spontaneous neurotransmitter release in *AKAP11*-deficient neuron

To investigate whether *AKAP11*-deficiency affects synaptic function, we performed electrophysiological recordings to measure miniature excitatory postsynaptic currents (mEPSCs) in *AKAP11*-cKO mice under the presence of tetrodotoxin (TTX) and GABAA receptor blocker picrotoxin (PTX). Given the critical role of the PFC in SCZ and BD pathophysiology^64–68^, we investigated neuronal transmission in layer 5/6 PFC of *Akap11*-cKO mice (Fig. 7a). Compared to the controls, *Akap11*-cKO neurons exhibited a significant decrease in mEPSC frequency and amplitude (Fig. 7b,c), indicating impaired excitatory synaptic transmission. To further investigate the underlying mechanisms, we also analyzed mEPSC kinetics and found no significant differences in rise time, decay, or half-width between groups (Extended Data Fig. 7a). However, both peak amplitude and area under the curve were significantly decreased in *Akap11*-cKO mice (Fig. 7d), suggesting diminished AMPA receptor-mediated synaptic responses.

**Fig. 7:**
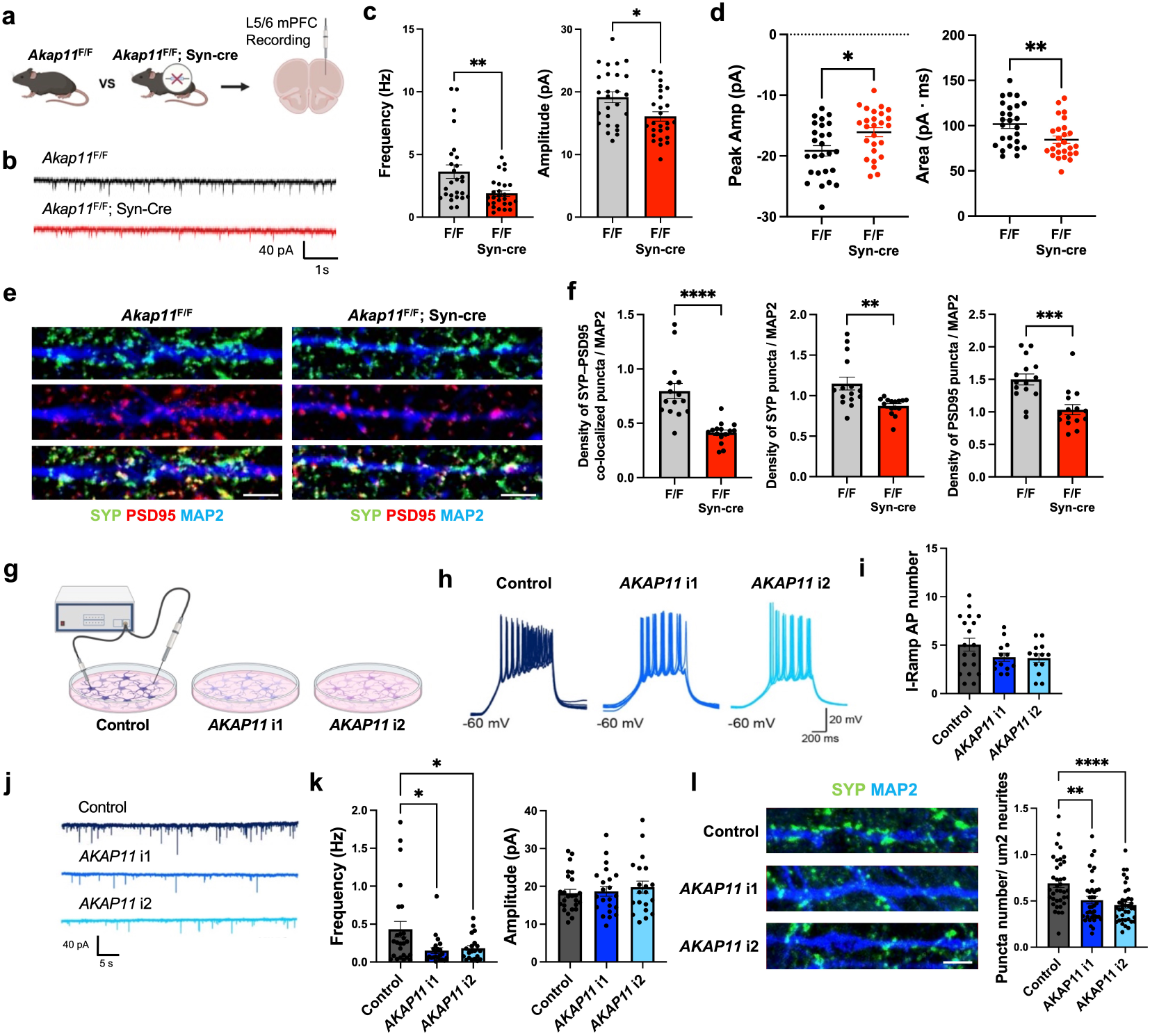
Electrophysiology analysis of *Akap11*-cKO mouse neurons and human *AKAP11*-KD iNeurons. **a**, Schematic images illustrating the electrophysiology experiments in *Akap11^F^*^/F^ and *Akap11*^F/F^; Syn-cre brain slices. **b**, Representative traces of mEPSC in the PFC of *Akap11^F^*^/F^ and *Akap11*^F/F^; Syn-cre mouse brains. **c**, Quantification of mEPSC frequency (left) and amplitude (right) in the PFC of *Akap11^F/F^* and *Akap11*^F/F^; Syn-cre brain slices. Data are presented as mean ± SEM (n>26 neurons, 3 independent replicates). Statistical significance (*p<0.05 and **p<0.01) was performed using an unpaired *t*-test. **d**, Quantification of mEPSC peak amplitude (left) and total charge transfer (area, right) in the PFC of *Akap11^F/F^* and *Akap11*^F/F^; Syn-cre brain slices. Data are presented as mean ± SEM (n>26 neurons, 3 independent replicates). Statistical significance (*p<0.05 and **p<0.01) was performed using an unpaired *t*-test. **e,f,** IF imaging of the primary cortical neurons from *Akap11^F^*^/F^ and *Akap11*^F/F^; Syn-cre mice, stained with anti-Synaptophysin, anti-PSD95, and anti-MAP2. Scale bars, 5μm. The bar graph shows quantification of SYP-PSD co-localized puncta, SYP puncta, and PSD95 puncta number normalized to MAP2. Data are presented as mean ± SEM (n>15 neurons, 3 independent replicates). Statistical analysis was performed using one-way ANOVA. **g**, Schematic illustration of the electrophysiology experiments in human *AKAP11*-KD iNeurons. **h,i,** Representative traces and quantification of induced action potentials (APs) number induced by a ramp depolarization protocol. Data are shown in mean ± SEM (n>20 neurons, 3 independent replicates). **j,k**, Representative traces (**h**) and quantification of mEPSC (**i**) (with 1uM TTX and 50uM PTX) frequency (left) and amplitude (right) in control and two clones of *AKAP11*-KD human iNeurons. Data are shown in mean ± SEM (n>20 neurons, 3 independent replicates). Statistical significance (*p<0.05) was evaluated with one-way ANOVA. **l**, IF imaging and quantification of the *AKAP11*-KD or Control iNeurons, stained with anti-Synaptophysin and anti-MAP2. Scale bars, 2μm. The bar graph shows quantification of puncta number per MAP2-positive neurite. Data are presented as mean ± SEM, (n > 37 per genotype). Statistical analysis was performed using one-way ANOVA.

To test whether the reduced mEPSC frequency was due to a loss of synapses, we performed IF staining in primary neurons. We found a significant reduction in the density of synaptophysin (SYP), PSD95, and co-localized SYP-PSD95 puncta in both *Akap11*-cKO and *Akap11*-wKO neurons compared to the control (Fig. 7e,f and Extended Data Fig. 7b), indicating a reduction in the number of mature excitatory synapse, which could explain the reduced mEPSC frequency.

Next, we investigated whether *AKAP11*-KD affects synaptic transmission in human iNeurons. To quantify changes in excitatory synaptic function, we recorded miniature excitatory postsynaptic currents (mEPSCs) under the same condition as in *Akap11*-cKO mice. Our analysis revealed a large decrease (∼3 fold) in the frequency of mEPSCs in *AKAP11*-KD human iNeurons, while the amplitude remained unaffected (Fig. 7g-k). Consistent with this, IF staining revealed a decrease in SYP-labeled signal density in *AKAP11*-KD iNeurons (Fig. 7l), phenocopying those in *Akap11*-cKO neurons, indicating disrupted synaptic vesicle protein homeostasis and impaired presynaptic transmission caused by AKAP11 deficiency.

## Discussion

Our current study reports a comprehensive analysis of autophagy receptor AKAP11’s cellular pathways and functions by employing multi-omics approaches, cell biology study, and electrophysiology analysis. Our data delineate a landscape of AKAP11-regulated proteome, kinome, transcriptome, and interactome, which reveals a central role of AKAP11 in coupling the type I PKA complex regulation to synaptic transmission in both human neurons and mouse models. Our study identifies a novel function of AKAP11 in autophagy by interacting with PKA-RI adaptors SPHKAP and VAPA/B. Given the genetic evidence linking *AKAP11*-coding variants to the shared risk of BD and SCZ^2,3,65^, our studies shed a light on the molecular and cellular mechanisms underlying the psychiatric diseases.

Our study demonstrates a pivotal role of AKAP11 in controlling homeostasis and activity of the type I PKA kinase and GSK3α/β in neurons. Multiple lines of evidence from our data reveal that *AKAP11*-deficiency leads to a disruption of homeostasis of PKA activity in neurons. Firstly, the proteomics study of *AKAP11*-KO neurons reveals the increase of regulatory subunits RIα/β and multiple inhibitors of PKA activity, including ADRA2A, PDE4B, PDE3A, and PKIB (Fig. 1b,e and Extended Data Fig. 1b); secondly, phosphoproteomics analysis identifies altered abundancy of predicted PKA phosphorylation sites (Fig. 3c); thirdly, the direct measurement of PKA activity showed a decrease of PKA activity in cytosol but an increase in synaptosome fractions of the brains of *Akap11*-cKO mice (Fig. 3d,e,f). The compartment-specific changes of the PKA activity in *AKAP11*-depleted neurons are not surprising, considering the functions of various AKAP in adapting PKA complex and regulating PKA activity in specific cellular compartments^69^. Interestingly, AKAP11 is highly enriched in cytosol and rarely detectable in synaptosomes fraction (Fig. 3d), consistent with the IF images of AKAP11 localization in the soma (Fig. 5b), whereas PKA-RIα/β and PKA-C are abundantly distributed in both cytosol and synaptosomes (Fig. 3d). The above data suggests the action of AKAP11 occurs primarily in the cytosol or soma, while the loss of AKAP11 impairs the selective autophagic degradation of PKA-RIα/β and PKA-C in the cytosol, resulting in an overflow of PKA-RIα/β and PKA-C proteins in multiple compartments including synaptosomes (Fig. 3d). However, the increase of RΙβ is much robust in the cytosol (6.41-fold) than in synaptosome (1.47-fold); in contrast, the PKA-C shows a stronger accumulation in the synaptosome (3.09-fold) than in the cytosolic (1.52-fold) (Fig. 3d,e). The observation explains the differential changes of PKA activity in the cytosol vs. synaptosomes.

Moreover, our data from phosphoproteome and experimental validation demonstrated a role for AKAP11 in regulating GSK3α/β kinase activity, which is also implicated in psychiatric disease^70–72^. AKAP11 directly binds GSK3α/β^5^; given GSK3α/β as substrates of AKT1^5,57,73^, the loss of AKAP11 may expose GSK3α/β protein and render the increase of phosphorylation of GSK3α/β by the elevated AKT1 activity (Fig. 4b-e). Therefore, it remains possible that the binding of AKAP11 prevents GSK3α/β from the inhibitory phosphorylation by AKT1, maintaining the active form of GSK3α/β. Interestingly, PKA was also shown to phosphorylate GSK3α/β at the same sites as by AKT1^6,57,74^. While the identification of the exact kinase responsible for GSK3α/β phosphorylation awaits future investigation, our data reveals a novel function and insight for AKAP11 in regulating GSK3α/β kinase activity through their direct binding. Further, our study suggests that AKAP11 level affects kinase activities beyond PKA and GSK3. But the predicted, extensive interactions among PKA, GSK3α/β, and those detected kinases (Fig. 4a and Extended Data Fig. 5b,c) raises a question that the changes of other kinase activities in *AKAP11*-deficient neurons could be a consequence of altered PKA and GSK3α/β activities.

Our study shows an important finding that AKAP11 couples the regulation of PKA signaling networks to synaptic transmission. The notion is supported by the evidence that a network module (WGCNA) shows a strong accordance of PKA-RI complex proteins with synaptic proteins, which are co-regulated by AKAP11(Fig. 2b,c,d,h). In *Akap11*-cKO neurons, the 399 phospho-proteins that are altered and predicted as PKA substrates display a significant enrichment in synaptic proteins (Fig. 3g,h), and the transcriptomics analysis reveals an enrichment of downregulated synapse related genes (Extended Data Fig. 2b,c). Furthermore, the depletion of AKAP11 proteins leads to impaired spontaneous synaptic transmission in both human and mouse neurons (Fig. 7c,d,k), demonstrating that an intact AKAP11 is critical for maintaining proper synaptic function through controlling type I PKA kinase and GSK3α/β activities. While these findings support a role for AKAP11 in regulating synaptic function via PKA, the mechanistic link between dysregulated PKA activity and synaptic impairment remains to be clarified. Additionally, the altered activities of other kinases such as GSK3α/β and AKT1 observed in AKAP11-deficient neurons may also contribute to synaptic dysfunction, providing a valuable direction for future investigation.

Through AKAP11-interactome analysis we uncovered the interactions between AKAP11and SPHKAP as well as VAP proteins in neurons. SPHKAP interacts with RIα/β and plays a role in adapting type I PKA to ER-PM junction through binding VAP proteins, which are ER-resident proteins^62,75,76^ and important for autophagosome biogenesis^58,59,77^. The interaction and colocalization of AKAP11 with VAP proteins therefore suggest the recruitment of AKAP11 to ER through VAP, which is involved in autophagosome biogenesis ^59,62,77^. Our study shows that AKAP11 is not required for the function of SPHKAP/VAP proteins in adapting PKA-RI to ER, but instead mediates SPHKAP degradation through autophagy, coinciding with the autophagic degradation of RIα/β (Fig. 6a,d,e,f). Together, our study provides an insight of AKAP11-mediated degradation of PKA-RI complex through autophagy and supports a model where AKAP11-SPHKAP interaction co-adapt PKA-RI complex to ER for degradation, while AKAP11-VAP interaction facilitates autophagosome biogenesis by recruiting ATG proteins to ER^58,59^. Thus, the AKAP11-SPHKAP-VAP complex offers an assembly platform to adapt type I PKA complex and recruit autophagy machinery at ER or ER-PM junction, leading to the selective degradation of PKA-RI complex. Such a coordinated action enables a precision control of PKA network homeostasis and compartmental activity coupled with calcium signaling, which dictates the synaptic transmission in neurons (Fig. 8a) ^60,61^.

**Fig. 8:**
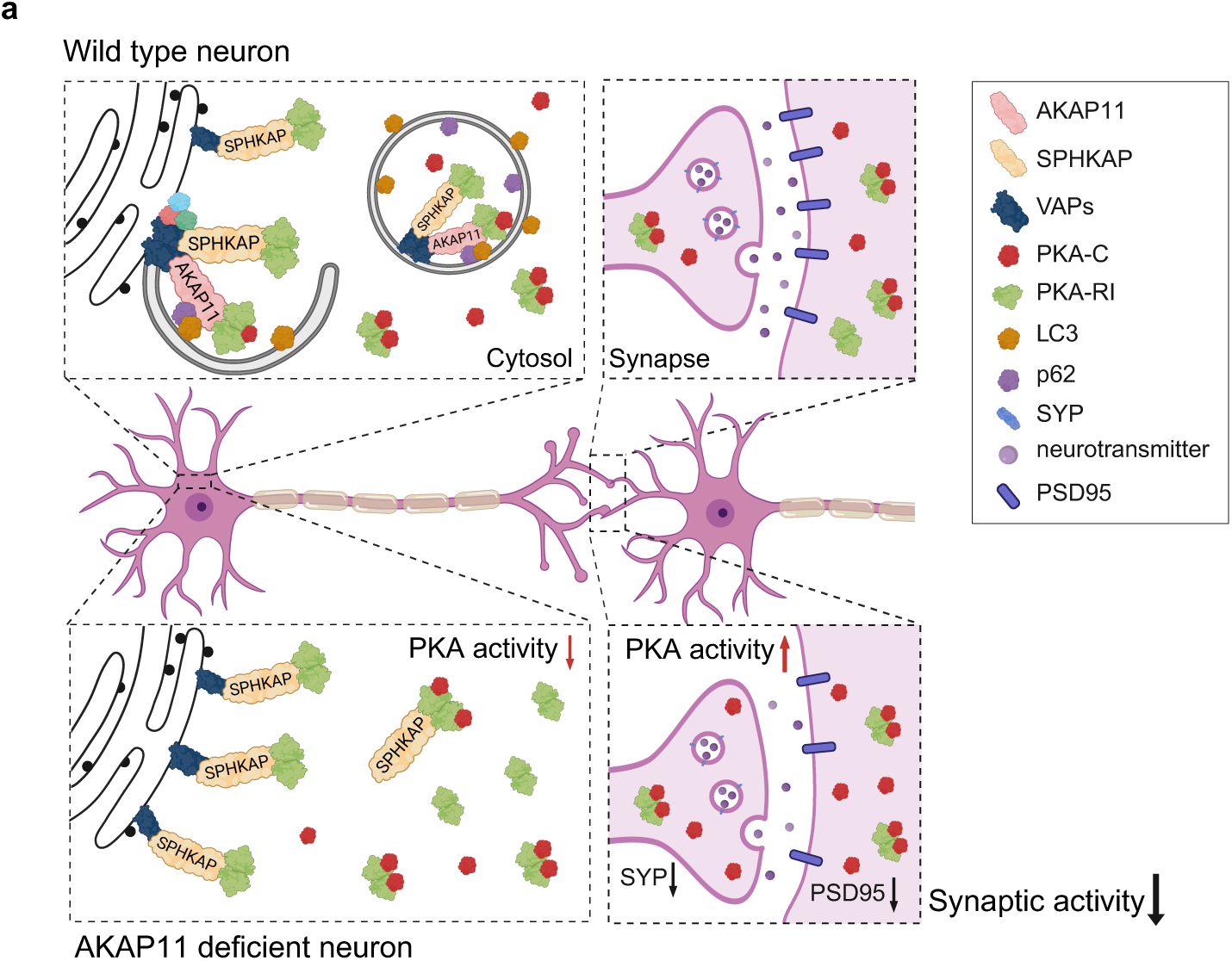
Graphical summary of AKAP11-regulated PKA signaling and synaptic function. **a**, In WT neurons, AKAP11 interacts with SPHKAP and VAPs at the ER to facilitate autophagic degradation of the PKA-RI complex, thereby maintaining compartment-specific PKA activity and normal expression of synaptic markers such as SYP and PSD95. In AKAP11-deficient neurons, impaired degradation leads to accumulation of PKA-RI and PKA-C, resulting in reduced cytosolic and increased synaptic PKA activity. This dysregulation is associated with decreased synaptic marker levels and impaired synaptic activity, highlighting AKAP11’s role in compartment-specific PKA regulation and synaptic function.

Although both mouse and human iPSC neuron models showed reduced mEPSC frequency following AKAP11 loss, only the mouse model exhibited decreased amplitude. This suggests that AKAP11-deficiency impairs presynaptic function or synapse number in human iNeurons, rather than postsynaptic strength. The differences may reflect species- or model-specific factors such as neuronal maturity, synaptic structure, or compensatory mechanisms. While our current study focuses on excitatory neurons, the potential contribution of inhibitory neurons to AKAP11-related phenotype remains an important and open question. Future studies will be necessary to investigate whether AKAP11 loss also impacts inhibitory synaptic function.

Finally, the DisGeNET annotation of the proteomics and phosphoproteomics data from *Akap11*-cKO neurons demonstrate a remarkable enrichment of psychiatric diseases including SCZ (top ranked), BD, and other mental disorders (Fig. 1i,3b). Given the link of protein-truncating variants of AKAP11 to the significant risk of SCZ and BD, the current characterization of the loss-of-function in autophagy receptor AKAP11 thus shed a light on a shared molecular mechanism underlying both SCZ and BD.

## Methods

### Animal models

All animal experiments were approved by the Institutional Animal Care and Use Committee (IACUC) of the Icahn School of Medicine at Mount Sinai and conducted in accordance with relevant ethical guidelines. Mice were housed in social cages under a 12-hour light/dark cycle with ad libitum access to food and water, and their food and water intake were monitored daily.

***Akap11* whole-body knockout (wKO) mice** (B6.Cg-Akap11<tm1.3Jsco/J>, strain #028921, discontinued) were obtained from The Jackson Laboratory. These mice were generated with Floxed exons 6 and 7 in *Akap11* allele. However, the mice suffered germline deletion of exons 6 and 7 without crossing Cre mice for unknown reason. Thus, the mutant mice were used as constitutive KO.

***Akap11* conditional knockout (cKO) mice** were generated with exon 6 flanked by loxP sites. The original mice were first generated using FLP recombinase (C57BL/6N-*A^tm1Brd^AKAP11^tm^*^1*a(EUCOMM)Hmgu*^/J-Mmucd, Stock number 046552-UCD, MMRRC) to remove the lacZ and Neo cassette, producing floxed (*Akap11*^flox/flox^) mice. These floxed mice were then crossed with Synapsin-Cre mice to generate *Akap11*-cKO mice, which were used to investigate the neuron-specific functions of *Akap11*.

***Akap11*-eGFP knock-in mice** were generated by fusing *Akap11* with eGFP and inserting it into the ROSA26 locus of B6.129S4-*Gt(ROSA)26^Sortm2(FLP*)Sor^*/J (strain #012930, Jackson lab) mice using a CRISPR/Cas9-based knock-in (KI) strategy, resulting in the targeted allele. These mice were then crossed with Synapsin-Cre mice to generate AKAP11-eGFP expressing mice for pan-neuronal expression. The strategy is based on the transcript NM_016248.4 and the design for the Cre reporter strains follows Srinivas et al. (2001) (BMC Dev Biol, 1:4). These mice were used to study the expression and localization of Akap11 in neurons.

### Primary neuronal culture

Mouse cortices from E16-18 embryos of pregnant females were dissected, and tissues were dissociated by incubating in 0.25% trypsin for 30 minutes at 37°C. The dissociated cells were plated on poly-L-lysin-coated coverslips in cell plates with growth medium composed of B27, Gluta-max, penicillin-streptomycin, and Neurobasal. After 18 days in vitro (DIV18), the cultures were fixed and prepared for immunostaining.

### Differentiation of iNeuron from human PSCs

Human PSCs were maintained and differentiated into glutamatergic neurons using Ngn2 overexpression system. PSCs were dissociated and plated in N2 medium supplemented with 50nM Chroman1, followed by transduction with FUW-TetO-Ngn2-P2A-puromycin and FUW-rtTA lentiviruses. Doxycycline was added to induce Ngn2 expression, and puromycin selection was applied to enrich transduced cells. Neurons were plated with mouse glial cells on Geltrex-coated plates and maintained in Neurobasal A medium with supplements, as previously described^10^. Mature neurons were used for experiments 5 weeks after co-culture initiation.

### Immunocytochemistry

Cells were fixed in 4% paraformaldehyde (PFA) in PBS for 20 minutes at room temperature and then washed three times with PBS. Permeabilization was performed with 0.3% Triton X-100 for 15 minutes and blocked with blocking solution (PBS containing 5% goat serum) for 1 hour at room temperature. Primary antibodies, Rabbit anti-AKAP11 (LS-Bio, LS-C374339), Rabbit anti-MAP2 (Abcam, ab5392), Sheep anti-RIβ (R&D, AF4177), Mouse anti-SPHKAP^60^, Guinea pig anti-Synaptophysin (Synaptic Systems, 101004), Guinea pig anti-p62 (PROGEN, GP62-C), diluted in blocking solution (PBS containing 5% goat serum and 0.2% Triton X-100), with were incubated overnight at 4°C. After washing three times with PBS, secondary antibodies, diluted in PBS, were incubated for 2 hours at room temperature. Images were captured using an LSM 900 confocal microscope, with 10X, 20X, 40X, 63X objectives and the images were processed using ImageJ.

### Immunohistochemistry

Mouse brains (2-5 months old) were perfused with 4% PFA, fixed overnight, and 30% sucrose for 2 days at 4°C. After sucrose removal, the brains were embedded in OCT compounds for cryosectioning and incubated at −80°C. The blocks were transferred to the cryostat 30 minutes before sectioning and incubated. The blocks were cut into 30µm thick sections at −20°C. The sections were placed in the 24-well plate with PBS to remove the OCT compound for 1hour at room temperature. Permeabilization was performed with 0.3% Triton X-100 for 30 minutes and blocked with blocking solution (PBS containing 0.3% Triton X-100 and 10% goat serum) for 1-hour at room temperature. Primary antibodies, Rabbit anti-AKAP11 (LS-Bio, LS-C374339), Rabbit anti-RIα (Cell signaling, 5675S), Sheep anti-RIβ (R&D, AF4177), Mouse anti-SPHKAP^60^, Guinea pig anti-Synaptophysin (Synaptic Systems, 101004), Rabbit anti-VAPA (Proteintech, 15275-1-AP), Rabbit anti-VAPB (Proteintech, 14477-1-AP), Guinea pig anti-p62 (PROGEN, GP62-C), diluted in blocking solution (PBS containing 5% goat serum and 0.2% Triton X-100), were incubated overnight at 4°C. After washing three times with PBS, secondary antibodies diluted in PBS containing 0.03% Triton X-100, were incubated for 2 hours at room temperature. Images were captured using an LSM 900 confocal microscope, with 10X, 20X, 40X, and 63X objectives using Z-stack and tile scan tools and analyzed with Image J.

### RNA extraction and Bulk RNA-seq DE analysis

Mouse brains, excluding the olfactory bulb and cerebellum, were transferred into RNase-free RINO tubes (Next Advance, Cat# PILR1-RNA) containing 1ml of RLT buffer from RNeasy Mini Kit (Qiagen). The tissues were homogenized using a Bullet Blender (Next Advance). After a brief centrifugation, the homogenates were diluted with additional RLT buffer. Total RNA was isolated by adding 70% ethanol to the homogenates and following the manufacturer’s protocol (RNeasy Mini Kit, Qiagen, Cat# 74016). All the RNA libraries were made and sequenced in Experimental Department in Novogene, USA, Sacramento, CA. Sequencing adapters were removed using Trimmomatic^78^ and the resulting reads were aligned to the GRCm39 primary assembly using HISAT2^79^. Gene counts for all samples were aggregated into a count matrix, which was subsequently used as input for DESeq2^80^ for statistical analysis. Principal component analysis (PCA) of the normalized, variance-stabilized count matrix revealed no outliers. Log2 fold changes and differential expression analysis were estimated using the DESeq function. A false discovery rate (FDR) threshold of 0.05 was applied to identify significantly differentially expressed genes.

### Western blot analysis of mouse brain lysate

After perfusion with PBS, whole brains were collected from mice, and the olfactory bulb and cerebellum were removed. The remaining brain tissue was homogenized using a Dounce homogenizer in lysis buffer containing 50mM Tris-HCl, pH 7.5, 150mM NaCl, 1% Triton X-100, and proteinase/phosphatase inhibitor. The homogenate was incubated on ice for 30 minutes and centrifuged at 1,200 x g for 10 minutes at 4°C. The supernatant was transferred to a new tube and centrifuged again at 15,000 x g for 20 minutes at 4°C. The final supernatant was collected on ice, and protein concentrations were measured by the Pierce BCA Protein Assay Kit following the manufacturer’s protocol.

Equal amounts of total proteins were loaded on 4%-12% Bis-Tris gel for SDS-PAGE and transferred to a 0.45µm PVDF membrane. Membranes were blocked in LI-COR Blocking Buffer (LI-COR, 927-60001) for 1 hour at room temperature. Primary antibodies, Rabbit anti-AKAP11 (LS-Bio, LS-C374339), Rabbit anti-RIα (Cell signaling, 5675S), Sheep anti-RIβ (R&D, AF4177), Mouse anti-RIIα (BD, 612242), Rabbit anti-PKACα (Cell signaling, 4782S), Rabbit anti-p-PKACα^T197^ (Cell signaling, 5661S), Mouse anti-β-actin (Cell signaling, 3700S), Rabbit anti-ADRA2A (Proteintech, 14266-1-AP), Rabbit anti-SPHKAP (Thermo, PA5-27581), Rabbit anti-Synaptophysin (Thermo, MA5-14532), Rabbit anti-GSK3α/β (Cell signaling, 5676S), Rabbit anti-p-GSK3α/β (S21/S9) (Cell signaling, 9331S), Rabbit anti-p-GluA1^S845^ (Cell signaling, 8084), Rabbit anti-GluA1 (Cell signaling, 13185), Rabbit anti-Akt1 (Cell signaling, 4691S), Rabbit anti-p-Akt1^S473^ (Cell signaling, 4060S), Chicken anti-GFP (Invitrogen, A10262), Rabbit anti-VAPA (Proteintech, 15275-1-AP), Rabbit anti-VAPB (Proteintech, 14477-1-AP), Guinea pig anti-p62 (PROGEN, GP62-C), Rabbit anti-LC3A/B (Cell signaling, 12742S), were diluted in blocking buffer with 0.2% Tween-20 and incubated overnight at 4°C. Secondary antibodies were diluted in blocking buffer with 0.2% Tween-20 and 0.01% SDS, followed by a 1-hour incubation at room temperature. Membranes were imaged using LI-COR imaging system and processed with ImageJ software.

### Immunoprecipitation

Brain tissue, excluding the olfactory bulb and cerebellum, was homogenized in sucrose buffer containing 250mM sucrose, 20mM Tris-HCl (pH 7.5), 0.5mM MgCl2, 0.5mM CaCl2, and protease/phosphatase inhibitors using a Dounce homogenizer. The homogenate was centrifuged at 1,000 x g for 10 minutes at 4°C. The supernatant was transfer to a prechilled tube and an equal volume of 2X RIPA buffer (40mM Tris-HCl, pH 7.5, 300mM NaCl, 2% NP-40, 1% sodium deoxycholate, and protease/phosphatase inhibitors) was added. The mixture was incubated at 4°C for 1 hour with rotation and then centrifuged at 15,000 x g for 20 minutes at 4°C. The supernatant was collected, and an aliquot was reserved as input.

For GFP-immunoprecipitation, 1mg of protein lysate was incubated overnight at 4°C with GFP-trap® magnetic particles (Chromotek, Cat# gtdk) under rotation. Beads were washed with RIPA buffer and bound proteins were eluted using 50µl of 1X SDS buffer by heat shock at 70°C for 10 minutes, repeated twice. The eluted proteins were subjected to SDS-PAGE. Subsequent steps followed the Western blot protocol described above.

For HA-immunoprecipitation, 500µg of protein lysates were incubated overnight at 4°C with anti-HA antibody (Santa Cruz, sc-7392) and anti-IgG antibody (Santa Cruz, sc-2025) under rotation. The next day, the lysates were incubated with Dynabeads^TM^ Protein G (Thermo, 10003D) for 2 hours at 4°C. After washing the beads with RIPA buffer, bound proteins were eluted with 30µl of 2X SDS sample buffer by heating at 95°C for 5 minutes. The eluted proteins were analyzed by SDS-PAGE followed by western blotting as described above.

### Purification of synaptic fractions

Synaptic fractions were isolated following a previously established protocol^81^. Briefly, mice brains were mechanically homogenized on ice using glass homogenizer (20 slow strokes) in a homogenization buffer containing 5mM HEPES (pH 7.4), 1mM MaCl_2_, 0.5mM CaCl_2_, and protease and phosphatase inhibitors. The homogenate was first centrifuged at 1,400 g for 10 min at 4°C, and the resulting supernatant was further centrifuged at 13,800 g for 10 min at 4°C to obtain the pellet. The resulting supernatant was collected as the cytosolic fraction. The pellet was resuspended in 0.32M sucrose, 6mM Tris-HCl (pH 7.5) and carefully layered onto a discontinuous sucrose gradient (0.85M, 1M, and 1.2M sucrose, all in 6mM Tris-HCl, pH 7.5). Ultracentrifugation was performed at 82,500 g for 2 hours at 4°C, and the synaptic fraction, located at the interface between the 1M and 1.2M sucrose layers, was collected. To extract synaptic proteins, the collected synapse was mixed with an equal volume of 1% Triton X-100 (in 6mM Tris-HCl, pH 7.5), incubated on ice for 15min, and centrifuged at 32,800 g for 20 min at 4°C. The final pellet, representing the synaptic fraction, was resuspended in lysis buffer for downstream analysis.

### Measurement of PKA activity

To measure PKA activity, perfusion with PBS was performed prior to tissue collection. Brain lysates from *Akap11*-cKO mice (excluding the olfactory bulb and cerebellum) and cortex/hippocampus lysates from *Akap11*-wKO mice were analyzed using the PKA activity kit (EIAPKA, Invitrogen) according to the manufacturer’s instructions.

Briefly, PKA standards or diluted samples, along with reconstituted ATP, were added to a PKA substrate-coated 96-well plate. The plate was incubated at 30°C for 90 minutes while shaking. After incubation, a Donkey anti-Rabbit IgG HRP-conjugated antibody was added, followed by the phospho-PKA substrate antibody, as per the protocol. The plate was incubated at room temperature while shaking for 60 minutes. TMB substrate was then added, and the reaction proceeded for 30 minutes at room temperature. Finally, stop solution was added, and the absorbance at 450nm was measured using a microplate reader.

### Protein extraction and digestion for mouse brain and iNeurons

Proteins in the tissues of mouse brains or iPSC cells were extracted into the lysis buffer by beads beating in a Bullet Blender at a speed of 6 for 30 s with an interval of 10s at 4^0^C for 6 cycles. The lysis buffer contained 8 M urea, 0.5% sodium deoxycholate, phosphatase inhibitor cocktails (Roche) and 50 mM HEPES pH 8.5. The proteins were digested with LysC (200:1 protein: enzyme ratio by weight, Wako) in the presence of 10 mM DTT and 5% acetonitrile for 3 h, followed by a 4-fold dilution with 50 mM HEPES buffer pH 8.5 and overnight trypsin digestion (50:1 protein : enzyme by weight, Promega) at room temperature. The peptides were desalted on C18 SPE column and dried under vacuum.

### Phosphopeptides enrichment of *Akap11*-cKO mouse brain samples

The phosphopeptides enrichment was carried out on TiO2 beads as previously described^82^. Briefly, ∼ 300 µg peptides were dissolved in 30 μL of binding buffer (65% acetonitrile, 2% TFA, and 1 mM KH2PO4). TiO2 beads (∼1 mg) were washed twice with washing buffer (65% acetonitrile, 0.1% TFA), mixed with the peptide solution, and incubated with end-over-end rotation at room temperature for 20 min. The phosphopeptide-bound beads were collected by brief centrifugation, washed twice with 200 μL washing buffer, and transferred to a C18 StageTip (Thermo Fisher Scientific). The phosphopeptides were eluted from StageTip with elution buffer (15 μL, 15% NH_4_OH, 40% acetonitrile). The eluents were dried under vacuum and stored at −80^0^c before MS analysis.

### Liquid Chromatography-Tandem Mass Spectrometry (LC-MS/MS) Analysis

For MS analysis, the dried peptides were dissolved in 5% formic acid and separated on a reverse phase column (50 µm × 15 cm, 1.7 µm C18 resin from CoAnn Technology) interfaced with a - timsTOP SCP MS (Bruker Daltonics) using nano-Elute 2 liquid chromatography system. Peptides were eluted at 55 °C by 5−27% buffer B gradient in 45 min (buffer A, 0.1% formic acid in water; buffer B, 0.1%formic acid in acetonitrile, flow rate of 0.25 μL/min). The peptides were ionized by a CaptiveSpray nanoelectrospray ion source, introduced into the MS and analyzed by a diaPASEF approach. The mass range was set as 100−1700 m/z and the ion mobility range was 0.70−1.30 1/K0.

### DIA based protein and phosphopeptides identification and quantification

To detect and quantify peptides and proteins, the raw data were searched by Spectronauts (version 18) against a mouse database that contains 55,260 entries or human database with 81,791 entries. Fixed and variable modifications included carbamidomethyl, methionine oxidation and N-termini acetylation. Additional phosphorylation at serine/threonine/tyrosine (S/T/Y) was enabled as variable modification for phosphoproteomic analysis. Both peptide and protein FDRs were controlled under 1%. Differentially expressed proteins/peptides were determined using both FDR and log2 fold change (FC) calculated by the limma^83^ R package. The significance of differential expression was evaluated based upon statistical criteria, with FDR < 0.05 and Log2(FC) > 2SD, or *P*-value < 0.05 and Log2(FC) > 2SD/1.5SD, followed by FDR estimation through permutation (FDR<0.05). The SD of proteins was estimated by fitting to a Gaussian distribution to evaluate the magnitude of experimental variations.

### TMT based protein identification and quantification

The TMT based proteomics analysis was performed as described before^84^. Briefly, the digested peptides were resuspended in 50 mM HEPES (pH 8.5) for TMTpro labeling, then were equally mixed and fractionated by offline basic pH reverse phase LC. Each of these fractions was analyzed by the acidic pH reverse phase LC-MS/MS on Q Exactive HF Orbitrap mass spectrometer. For mass spectrometer settings, positive in mode and data-dependent acquisition were applied with one full MS scan followed by a 20 MS/MS scans. MS1 scans were collected at a resolution of 60,000, 1 × 10^6^ AGC and 50 ms maximal ion time; MS2 spectra were acquired at a resolution of 60,000, fixed first mass of 120 m/z, 410–1600 m/z, 1 × 10^5^ AGC, 110ms maximal ion time, and ∼15 s of dynamic exclusion. The TMT dataset was processed by the JUMP software suite. The raw files were searched against a mouse database with 59,423 entries downloaded from Swiss-Prot, TrEMBL, and UCSC databases. Main search parameters were set at precursor and product ion mass tolerance (± 15 ppm), full trypticity, maximal modification sites (n = 3), maximal missed cleavage (n = 2), static mass shift including carbamidomethyl modification (+57.02146 on Cys), TMT tags (+304.20714 on Lys and N-termini), and dynamic mass shift for oxidation (+15.99491 on Met). Peptide-spectrum matches (PSM) were filtered by mass accuracy, clustered by precursor ion charge, and the cutoffs of JUMP-based matching scores (J-score and ΔJn) to reduce FDR below 1% for proteins.

### Weighted Gene Correlation Network Analysis (WGCNA)

The analysis was carried out with the WGCNA^39^ R package. For the *Akap11*-cKO mouse brain proteomic data, we used the blockwiseModules function to generate the network, with the settings soft-threshold power = 16, minimum module size = 30, network type = ‘‘signed’’, TOMType = “signed”, randomSeed = 1234, pamRespectsDendro = FALSE, maxBlockSize = 20,000, and merge cut height = 0.25. The partitioning around medoids-like (PAM-like) stages of module detection was disabled to ensure that module assignments adhered strictly to the hierarchical clustering dendrogram (pamRespectsDendro = FALSE). The soft-threshold power of 16 was chosen based on the fitted signed scale-free topology model (R2) approaching an asymptote of 0.85 at that threshold. Furthermore, the mean connectivity at that power was less than 100. The maximum block size was set at 20,000 to complete all clustering within a single block. To get a more robust result, combined our *Akap11*-cKO mouse brain proteomic dataset with our previous work^10^. While for the *AKAP11*-KD iNeuron proteomic data, the settings were: soft-threshold power = 18, minimum module size = 80, network type = ‘‘signed’’, TOMType = “signed”, randomSeed = 1234, pamRespectsDendro = FALSE, maxBlockSize = 20,000. We also combined our *AKAP11*-KD iNeuron proteomic data with our previous work^10^.

### Protein-Protein Interaction (PPI) Network Analysis

The protein-protein interaction network analysis was performed using the STRING^41^ plugin in CytoScape^85^, with the fold changes and FDR/*P*-values obtained from the differentially expressed proteins analysis. Briefly, for the Akap11-cKO mouse brain proteomic data in ME15, we analyzed the PPI network among these proteins using STRING. Subsequently, we adjusted the node colors and sizes based on the fold changes and FDR obtained from differentially expressed protein analysis presented in Fig. 1b. We performed a similar analysis with the enriched proteins from the AKAP11 interactome data.

### Pathway enrichment analysis

GO term and GSEA were performed with the R Bioconductor package clusterProfiler^23^. Briefly, we first transfer the proteins symbols to Entrez IDs based on Bioconductor annotation data package (org.Hs.eg.db/org.Mm.eg.db) and used the clusterProfiler::enrichGO or clusterProfiler::gseGO with the pvalueCutoff = 0.05, qvalueCutoff = 0.05 to do pathway enrichment analysis. For human disease enrichment analysis, we utilized the ToppGene Suite^86^ in conjunction with the DisGeNET^38^ database to do enrichment analysis and generate the dot plot utilizing the ggplot2^87^ package. All the volcano plots were generated by EnhancedVolcano^88^ R package.

### Phosphoproteomic data analysis

The analysis of phosphoproteomic data was generally followed as outlined in the paper introducing PhosR^51^, which employs a multi-step framework comprising two major components: the first components is the kinaseSubstrateScore function, which evaluates a given phosphosite by integrating kinase recognition motifs and phosphoproteomic dynamics, and the second component is the kinaseSubstratePred function, which combines the scores from the first component for predicting kinase-substrate relationships using an adaptive sampling-based positive-unlabeled learning method^89^. Briefly, we first did the PCA analysis and deleted the outlier one sample in each group (3 Control vs. 3 *Akap11*_cKO) and modified the format of our phosphoproteomic data to conform to the input requirements for PhosR^51,53^. The general analysis workflow comprises the following steps: log transformation, imputation, differentially abundant phosphoprotein site analysis, identifying stably phosphorylated sites, normalization and batch correction of datasets, predicting kinase substrates, and constructing signaling networks (signalomes).

### Slice Electrophysiology

Electrophysiological recordings from acute prefrontal slices were performed essentially as described previously^90^. Briefly, slices were prepared from *Akap11*-cKO mice aged 2–3 months. Coronal brain slices (300μm thickness) were cut in a high-sucrose cutting solution containing (in mM): 85 NaCl, 75 sucrose, 2.5 KCl, 1.3 NaH₂PO₄, 24 NaHCO₃, 0.5 CaCl₂, 4 MgCl₂, and 25 D-glucose. Slices were equilibrated in artificial cerebrospinal fluid (ACSF) at 31 °C for 30 minutes, followed by an additional hour at room temperature. Slices were then transferred to a recording chamber containing ACSF with the following composition (in mM): 120 NaCl, 2.5 KCl, 1 NaH₂PO₄, 26.2 NaHCO₃, 2.5 CaCl₂, 1.3 MgSO₄·7H₂O, and 11 D-glucose (∼290 mOsm), supplemented with 50 μM picrotoxin and 0.5 μM TTX. Miniature excitatory postsynaptic currents (mEPSCs) from Layer 5–6 cells in the medial prefrontal cortex (mPFC) were recorded in whole-cell voltage-clamp mode (holding potential = −70 mV) using an internal solution with the following composition (in mM): 117 Cs-methanesulfonate, 15 CsCl, 8 NaCl, 10 TEA-Cl, 0.2 EGTA, 4 Na₂-ATP, 0.3 Na₂-GTP, and 10 HEPES, pH 7.3 adjusted with CsOH (∼300 mOsm). All recordings were acquired using Clampex 10 data acquisition software (Molecular Devices) and a Multiclamp 700B amplifier (Molecular Devices). Miniature events were manually identified using a 5pA threshold in the Clampfit software.

### Patch Clamp electrophysiology in human iN cells

Whole-cell patch-clamp electrophysiology for iNeurons cells was performed as described^91^. To examine intrinsic membrane properties and record action potentials, K-gluconate internal solution (in mM): 126 K-gluconate, 4 KCl, 10 HEPES, 0.05 EGTA, 4 ATP-magnesium, 0.3 GTP-sodium, and 10 phosphocreatine was used. Miniature postsynaptic currents were recorded at a holding potential of −70 mV in the presence of 1 μM tetrodotoxin (TTX) and 50 μM Picrotoxin (PTX). The Cs-based internal solution used for this purpose contained the following components (in mM): 40 CsCl, 3.5 KCl, 10 HEPES, 0.05 EGTA, 90 K-gluconate, 1.8 NaCl, 1.7 MgCl_2_, 2 ATP-magnesium, 0.4 GTP-sodium, and 10 phosphocreatine.

The external solution used during recordings consisted of (in mM): 140 NaCl, 5 KCl, 10 HEPES, 2 CaCl_2_, 2 MgCl_2_, and 10 Glucose, with a pH of 7.4. All recordings of cell culture experiments were performed at room temperature.

To ensure data quality, any cells exhibiting a change in series resistance (Rs) exceeding 20% during the recording were excluded from the analysis. Additionally, recordings with an access resistance greater than 25 MΩ were also excluded. The electrophysiological data were collected using pClamp 10.5 software (Molecular Devices) and the MultiClamp 700B amplifier (Molecular Devices). The detection of miniature excitatory postsynaptic currents (mEPSCs) was performed using the template search algorithm in Clampfit 10.5 software (Molecular Devices).

### Quantification and Statistical Analysis

Unless otherwise specified, all data are presented as the means of at least three biological replicates, each derived from a minimum of three independent experiments. Statistical significance between two groups was evaluated using an unpaired, two-tailed Student’s t-test. For comparisons involving multiple means, statistical significance was assessed using one-way analysis of variance (ANOVA) followed by Dunnett’s multiple comparisons test, conducted in GraphPad Prism (version 10.2.3; GraphPad Software, CA, USA). Data are reported as mean ± SEM, with specific details regarding statistical tests provided in the respective figure legends. Error bars represent the standard deviation of all replicates. Graphs and plots were generated using GraphPad Prism (version 10.2.3) or RStudio (version 2024.09.1).

## Data availability

The authors affirm that all supporting data for the study’s findings are readily available within the paper and its Supplementary Information files. For further inquiries regarding resources and reagents, please contact the lead contact, Zhenyu Yue, Nan Yang, Junmin Peng, and Jinye Dai. The source data accompanying this publication are also provided.

## Acknowledgements

We are grateful to Dr. Vierra for kindly providing the SPHKAP antibody. The schematics were created using BioRender. This work was supported by the NIH grants R01NS060123 (Z.Y., N.Y., J.P.), R01NS117590 and R21AG067570 (Z.Y. and N.Y.). Z. Pang’s laboratory is supported by R01MH131296, R01MH125528 and RM1MH133065. L.W. was supported by the New Jersey Governor’s Council for Medical Research and Treatment of Autism Postdoctoral Fellowship (CAUT24DFP).

## Author contributions

Zhenyu Yue conceived the study. You-Kyung Lee performed the sample preparation, immunoblot assay, GFP affinity purification, immunoflueoscence staining with *Akap11*-cKO, *Akap11*-wKO mouse brains, and *AKAP11*-deficiency neurons. Cong Xiao did the data analysis of the proteomic data from *Akap11*-cKO and *Akap11*-wKO mouse brains and *AKAP11*-KD iNeurons, the RNAseq data from *Akap11*-cKO, as well as the phosphoproteomic data from *Akap11*-cKO mouse brains. Xianting Li generated *AKAP11*-deficient mouse models and performed the interactome sample preparation, silver staining and GFP affinity purification using AKAP11-eGFP mouse brains. Xiaoting Zhou and Henry Kim assisted generating and analyzing of proteomics of iNeurons. Zhiping Wu and Meghan G. McReynolds performed the proteomics and phospho-proteomics analysis by MS. Le Wang did the electrophysiology analysis of *AKAP11*-KD iNeurons guided by Zhiping Pang. Jinye Dai did the electrophysiology analysis of *Akap11*-cKO mouse neruons. Junmin Peng guided the proteomics data analysis and data interpretation. Zhenyu Yue, You-Kyung Lee, Cong Xiao, Nan Yang, Junmin Peng drafted the manuscript.

## Supplementary Figures

**Extended Data Fig. 1:**
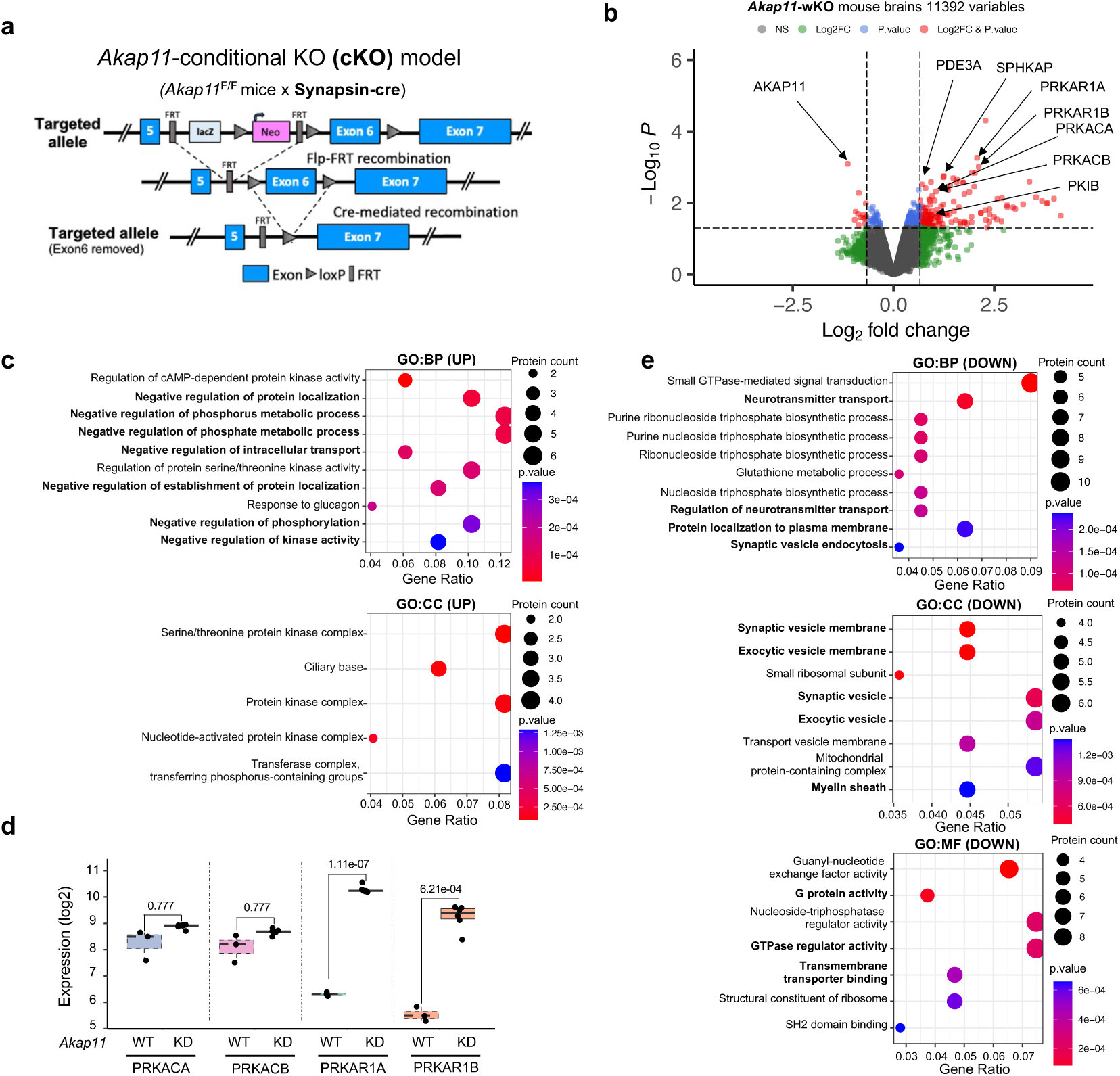
Intergrated proteomics analysis of *Akap11*-cKO and *Akap11*-wKO mouse brains, and *AKAP11*-KD human iNeurons. **a**, The strategy for generating the *AKAP11*-cKO mouse model. **b**, Volcano plot of the differentially expressed proteins (DEPs) in *Akap11*-wKO mouse brains. A positive score indicates enrichment, a negative score indicates depletion. The y axis represents statistical confidence (*P*-value) for each x axis point. Enriched proteins, defined by *P*-value < 0.05 and |Log2FC| > 2SD and followed by FDR estimation through permutation (FDR<0.05), are colored in red. **c**,**e**, Gene Ontology (GO) annotations of the DEPs in Fig. 1b, displaying the top 10 enriched pathways with *P*-value < 0.05. **d**, Enrichment levels for the proteins as shown in Fig. 1c with log2 transformed TMT intensities across AKAP11-eGFP and control mice.

**Extended Data Fig. 2:**
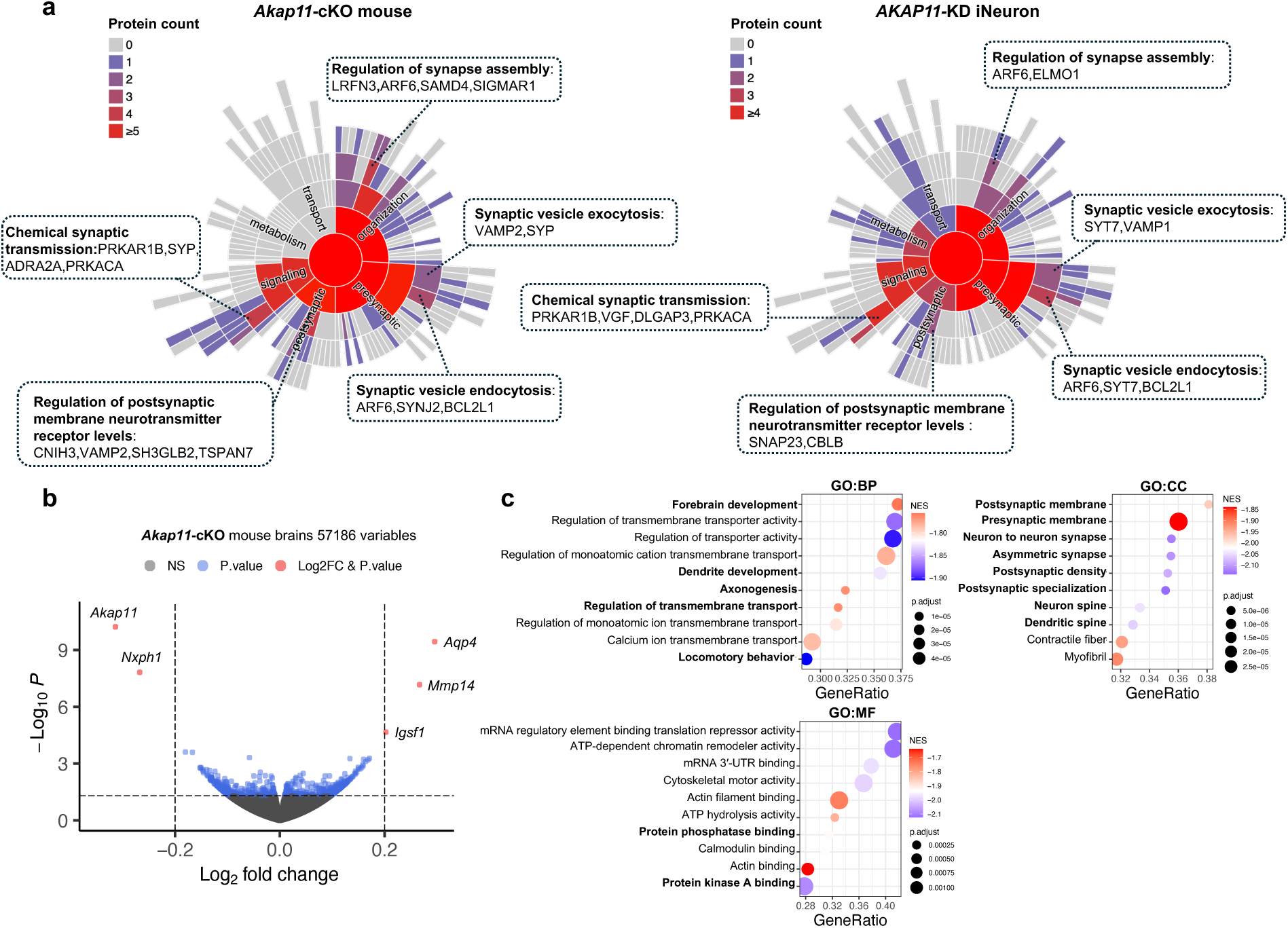
Synapse protein enrichment and transcriptomic analysis of brain lysate from *AKAP11*-deficient neurons. **a**, Sunburst plots depicting SynGO Biological Process pathways with a *P*-value < 0.05. For selected ontologies, representative DEPs in the ontology are displayed. **b**, Volcano plot of the differentially expressed genes (DEGs) in *Akap11*-cKO mouse brains. A positive score indicates enrichment, a negative score indicates depletion. The y axis represents statistical confidence (adjust *P*-value) for each x axis point. Enriched genes, defined by having adjusted *P*-value < 0.05 and Log2FC > 0.2, are colored in red. **c**, Gene set enrichment analysis (GSEA) of all protein Gene set enrichment analysis of all proteins displayed in **a**, showing the top 10 ontologies with a negative normalized enrichment score (NES) and adjusted *P*-value < 0.05.

**Extended Data Fig. 3:**
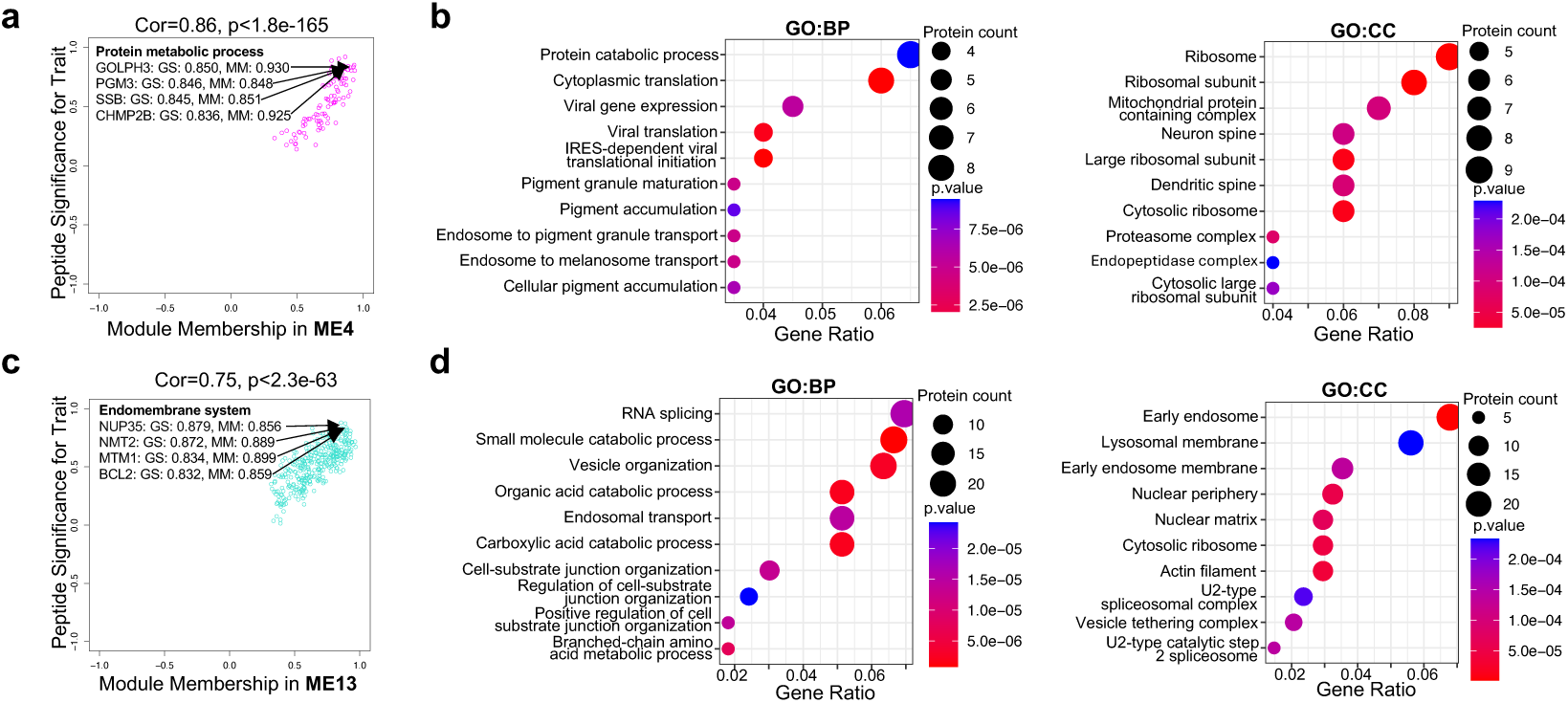
Identification of co-regulation of cellular pathways in positive corelated protein modules upon *AKAP11*-deficiency through WGCNA. **a**,**c**, Scatter plot of the module membership (X axis) and peptide significance (Y axis) in ME4 from *Akap11*-cKO mouse brains (**a**) or ME13 from *AKAP11*-KD from human iNeurons (**c**). **b**,**d**, Gene Ontology (GO) annotations of the proteins in ME4 from *Akap11*-cKO mouse brains (**b**) or ME13 from *AKAP11*-KD from human iNeurons (**d**), showing top 10 enriched pathways with *P*-value < 0.05.

**Extended Data Fig. 4:**
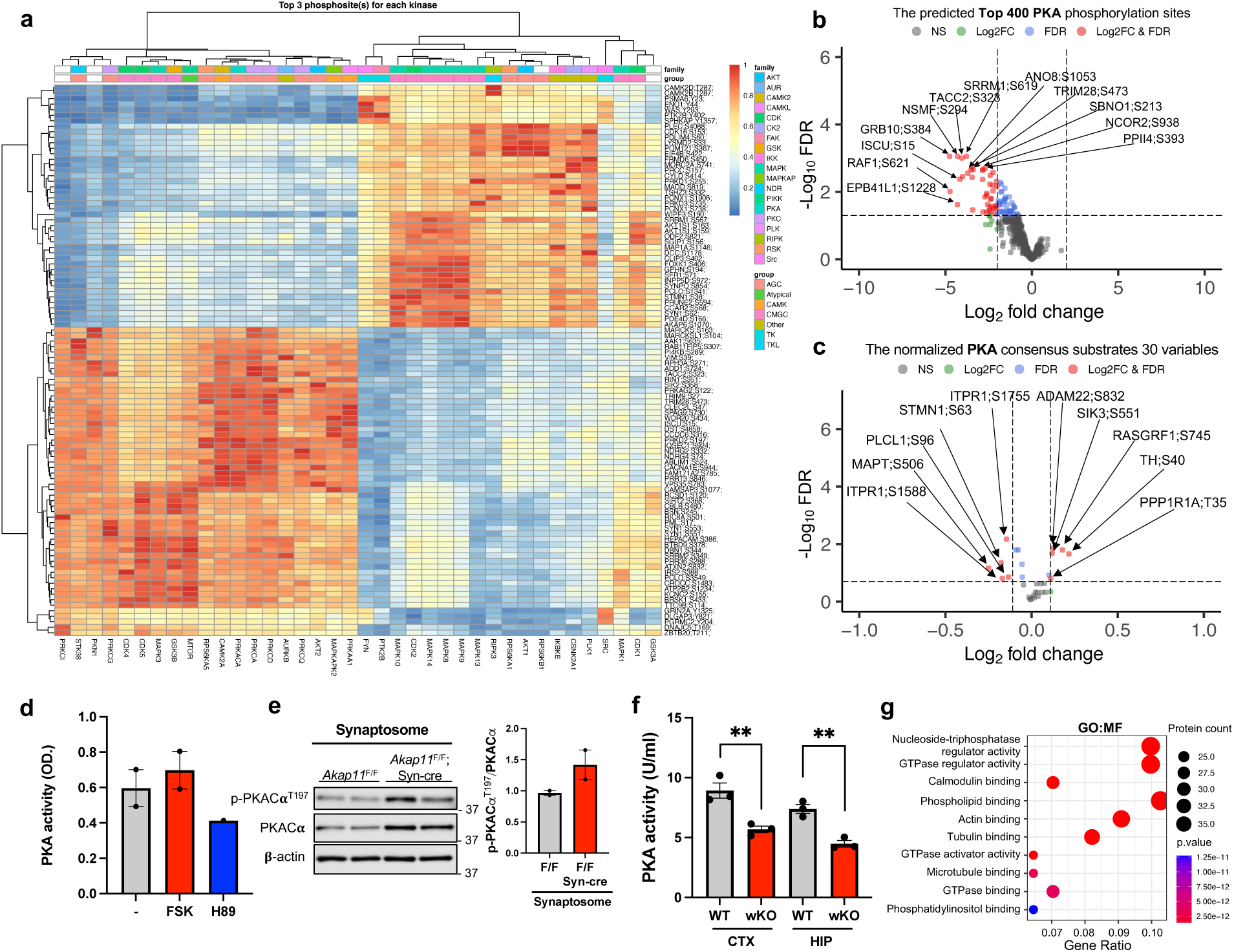
Exploration of altered kinase activities in *Akap11-*cKO brain lysates. **a**, Heatmap of the top three phosphoprotein sites for each kinase. The differentially abundant phosphoprotein sites (y axis) and the kinases (x axis) are plotted. **b**, Volcano plot of the top 400 predicted PKA phosphorylation sites detected by mass-spectrometry in *Akap11*-cKO mouse brains. A positive score indicates enrichment, a negative score indicates depletion. The y axis represents statistical confidence (adjusted *P*-value (FDR)) for each x axis point. Enriched proteins, defined by FDR < 0.05 and Log2FC > 2SD, are colored in red. **c**, Volcano plot of the PKA consensus substrates (normalized by the total protein level) detected by mass-spectrometry in *Akap11*-cKO mouse brains. A positive score indicates enrichment, a negative score indicates depletion. The y axis represents statistical confidence (adjusted *P*-value (FDR)) for each x axis point. Enriched proteins, defined by having FDR < 0.1 and Log2FC > SD, are colored in red. **d**, Bar graph of the results from the ELISA of PKA activity from primary neurons treated with 10μM forskolin (FSK) or 5μM H89 treatment for 30 minutes. Data are presented as mean ± SEM. **e**, Immunoblotting using antibodies against PKACα and p-PKACα^T197^ in synaptosome fraction and quantification of the results. Data are presented as mean ± SEM. Statistical analysis was performed using an unpaired *t*-test. **f,** Bar graph of the results from the ELISA of PKA activity in the cytosolic fraction of the cerebral cortex and hippocampus of *Akap11*-wKO mice. Brain tissues were homogenized in 50mM Tris-HCl (pH 7.5), 0.32M sucrose, 1mM MaCl_2_, 2.5mM CaCl_2_, and 1mM NaCl before analysis. Data are presented as mean ± SEM (n=3 per genotype). Statistical analysis was performed using an unpaired *t*-test. **g,** Gene Ontology (GO) annotations of the predicted PKA substrates in Fig. 3g, displaying the top 10 enriched pathways with *P*-value < 0.05.

**Extended Data Fig. 5.**
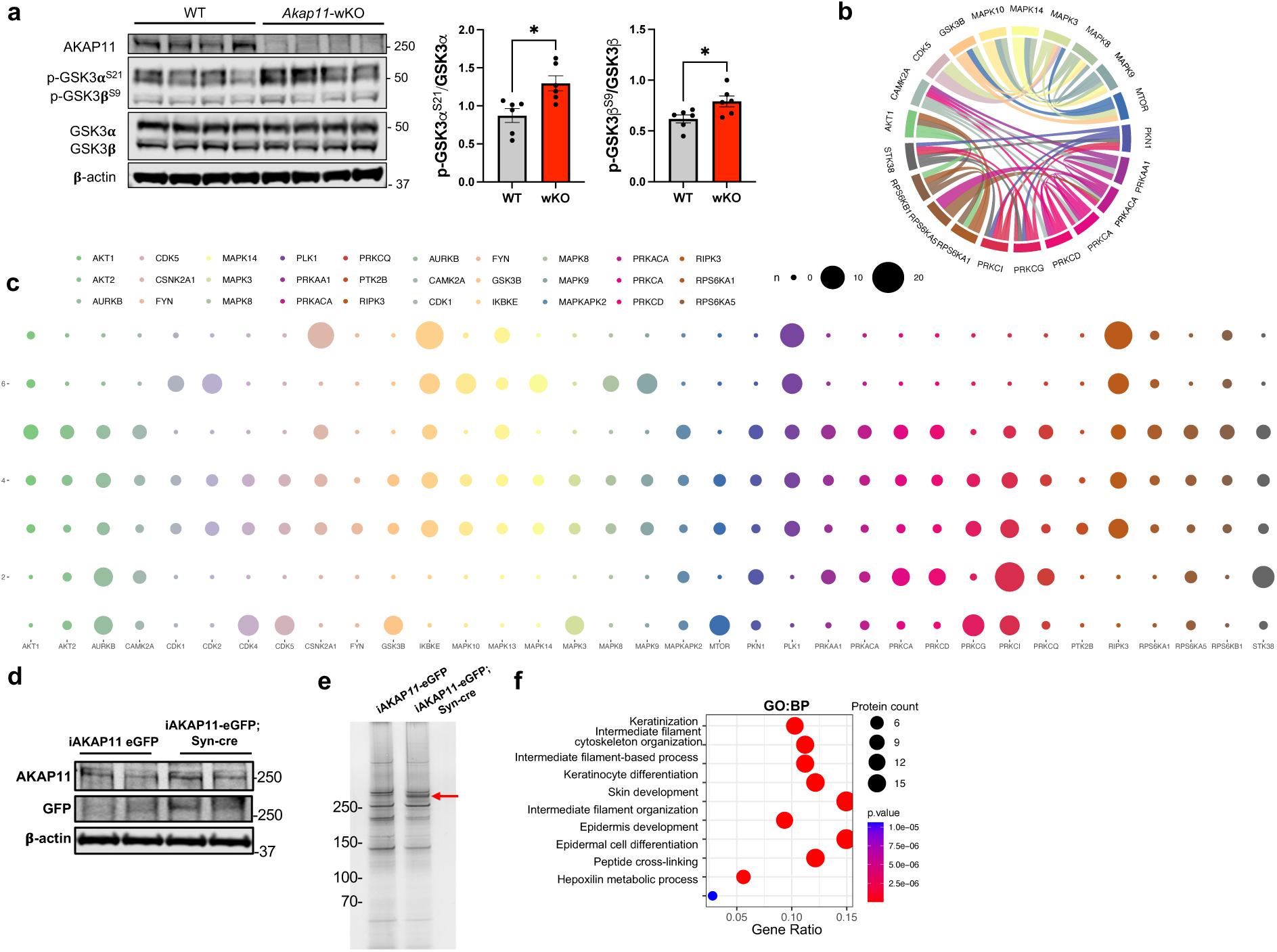
Exploration of kinase network and pattern. **a,** Immunoblotting using antibodies against AKAP11, GSK3α/β and p-GSK3α/β(S21/S9) and quantification of the blot results. Data are presented as mean ± SEM (n=6 per genotype). Statistical analysis was performed using an unpaired *t*-test. **b**, Kinase network analysis of all detected kinases in *Akap11*-cKO mouse brains. The nodes represent kinases; the edges refer to the *Pearson*’s correlation of kinase regulation between the two nodes. The wider the edge, the stronger the correlation. **c**, Balloon plot of the patterns for each kinase. The y-axis represents the distribution of the predicted substrates in seven protein modules for each kinase (x-axis). **d**, Immunoblotting using antibodies against AKAP11, GFP and β-actin. **e,** Silver stained SDS-PAGE for IP samples from cytoplasm of AKAP11-eGFP and AKAP11-eGFP; Syn-cre mouse brains. 10% of the IP sample was loaded from each IP sample. Arrow indicated IP-ed AKAP11-eGFP protein. **f**, Gene Ontology (GO) annotations of the proteins shown in Fig. 5f, displaying the top 10 enriched pathways with *P*-value < 0.05.

**Extended Data Fig. 6:**
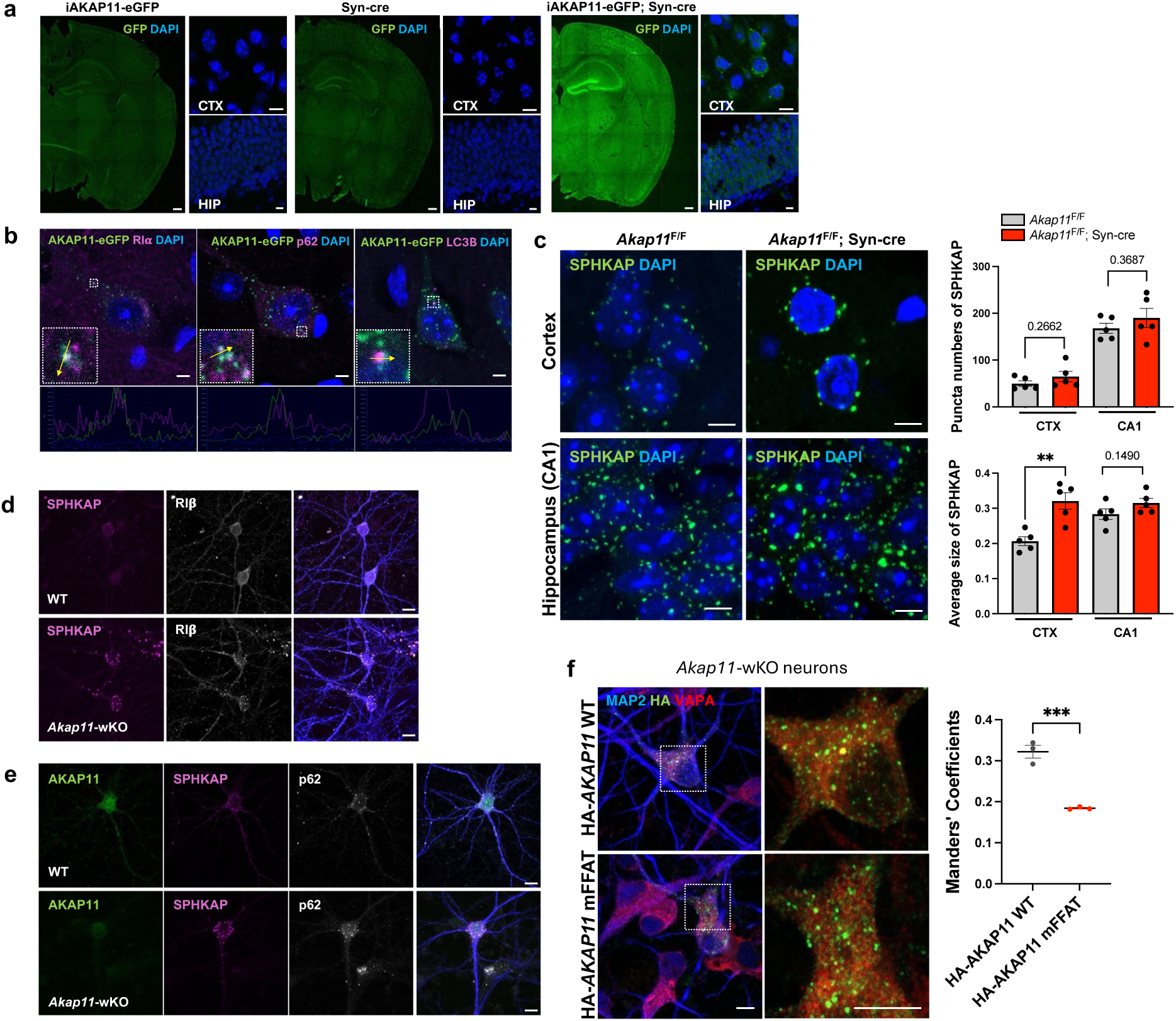
Characterization of SPHKAP in neurons from *Akap11*-cKO mouse brains. **a**, Immunofluorescence images of cortical sections from AKAP11-eGFP, Syn-cre, and AKAP11-eGFP; Syn-cre mice brain, showing AKAP11 expression in overall brain regions. Inset images provide magnified views of cortex and hippocampus. Scale bars, 200μm (main images); 10μm (inset images). **b**, Representative immunofluorescence images of AKAP11-eGFP transgenic mice brain slices stained with anti-RIα and anti-VAPB antibodies in the cortex, showing co-localization of these proteins. Scale bars, 5μm. **c**, Immunofluorescence images of *Akap11*^F/F^ and *Akap11*^F/F^ Syn-cre mice brain stained with anti-SPHKAP antibody. Scale bars, 5μm. Quantification of SPHKAP puncta numbers and average size using Analyze Particles in ImageJ. Data are presented as mean ± SEM. Statistical significance (**p<0.01) was performed using an unpaired t-test. **d**, Immunofluorescence images of primary neuron from WT and *Akap11*-wKO mice stained with anti-SPHKAP, anti-RIβ, and anti-MAP2 antibodies. Scale bars, 10μm. **e**, Immunofluorescence images of primary neuron from WT and *Akap11*-wKO mice stained with anti-AKAP11, anti-SPHKAP, anti-p62, anti-MAP2 antibodies. Scale bars, 10μm. **f,** Primary neurons from *Akap11*-wKO mice expressing HA-AKAP11 WT or mFFAT were stained with antibodies against HA, VAPA, and MAP2. The bar graph shows quantification of HA–VAPA co-localized signal using Manders’ coefficient. Data are presented as mean ± SEM (n > 3 per group). Statistical significance (***p<0.001) was performed using an unpaired *t*-test. Scale bars, 5 μm.

**Extended Data Fig. 7:**
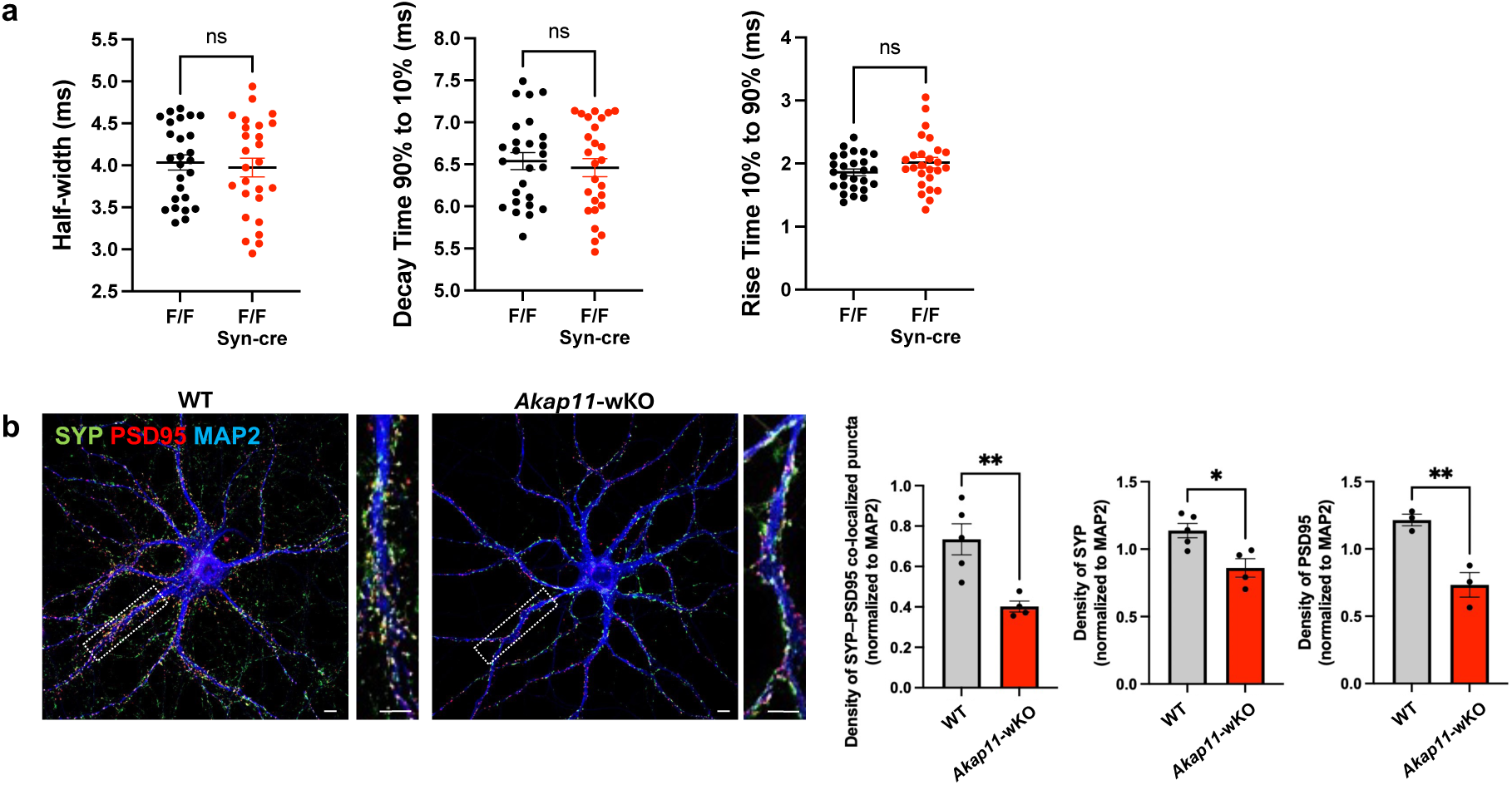
Impaired spontaneous neurotransmitter release in *AKAP11*-deficient neurons. **a**, Quantification of mEPSC kinetics in the PFC of *Akap11^F/F^* and *Akap11*^F/F^; Syn-cre brain slices. Kinetic parameters include half-width (duration at 50% of peak amplitude), decay time (time to decay from 90% to 10% of peak), and rise time (time to rise from 10% to 90% of the peak). Data are presented as mean ± SEM (n>26 neurons, 3 independent replicates). Statistical significance was performed using an unpaired *t*-test. **b,** Immunofluorescence imaging and quantification of the WT and *Akap11*-wKO stained with anti-Synaptophysin, anti-PSD95, and anti-MAP2. Scale bars, 5μm. The bar graph shows quantification of puncta number per μm^2^ neurite. Data are presented as mean ± SEM (n>4 per genotype). Statistical analysis was performed using an unpaired *t*-test (*p<0.05, **p<0.01, and ***p<0.001).

## REFERENCES

1. Palmer, D. S. et al. Exome sequencing in bipolar disorder identifies AKAP11 as a risk gene shared with schizophrenia. Nat. Genet. 54, 541–547 (2022).

2. Liu, D. et al. Schizophrenia risk conferred by rare protein-truncating variants is conserved across diverse human populations. Nat. Genet. 55, 369–376 (2023).

3. Singh, T. et al. Rare coding variants in ten genes confer substantial risk for schizophrenia. Nature 604, 509–516 (2022).

4. Lester, L. B., Coghlan, V. M., Nauert, B. & Scott, J. D. Cloning and Characterization of a Novel A-kinase Anchoring Protein. J. Biol. Chem. 271, 9460–9465 (1996).

5. Whiting, J. L. et al. Protein Kinase A Opposes the Phosphorylation-dependent Recruitment of Glycogen Synthase Kinase 3β to A-kinase Anchoring Protein 220. J. Biol. Chem. 290, 19445–19457 (2015).

6. Fang, X. et al. Phosphorylation and inactivation of glycogen synthase kinase 3 by protein kinase A. Proc. Natl. Acad. Sci. 97, 11960–11965 (2000).

7. Tanji, C. et al. A-Kinase Anchoring Protein AKAP220 Binds to Glycogen Synthase Kinase-3β (GSK-3β) and Mediates Protein Kinase A-dependent Inhibition of GSK-3β. J. Biol. Chem. 277, 36955–36961 (2002).

8. Wong, W. & Scott, J. D. AKAP signalling complexes: focal points in space and time. Nat. Rev. Mol. Cell Biol. 5, 959–970 (2004).

9. Deng, Z. et al. Selective autophagy of AKAP11 activates cAMP/PKA to fuel mitochondrial metabolism and tumor cell growth. Proc. Natl. Acad. Sci. 118, e2020215118 (2021).

10. Zhou, X. et al. Integrated proteomics reveals autophagy landscape and an autophagy receptor controlling PKA-RI complex homeostasis in neurons. Nat. Commun. 15, 3113 (2024).

11. Herzog, L. E. et al. Mouse mutants in schizophrenia risk genes GRIN2A and AKAP11 show EEG abnormalities in common with schizophrenia patients. Transl. Psychiatry 13, 92 (2023).

12. Aryal, S. et al. Deep proteomics identifies shared molecular pathway alterations in synapses of patients with schizophrenia and bipolar disorder and mouse model. Cell Rep. 42, 112497 (2023).

13. Kallergi, E. et al. Dendritic autophagy degrades postsynaptic proteins and is required for long-term synaptic depression in mice. Nat. Commun. 13, 680 (2022).

14. Ji, C., Tang, M., Zeidler, C., Höhfeld, J. & Johnson, G. V. BAG3 and SYNPO (synaptopodin) facilitate phospho-MAPT/Tau degradation via autophagy in neuronal processes. Autophagy 15, 1199–1213 (2019).

15. Chang, Y.-C., et al. Identification of secretory autophagy as a mechanism modulating activity-induced synaptic remodeling. Proc. Natl. Acad. Sci. 121, e2315958121 (2024).

16. Nikoletopoulou, V., Sidiropoulou, K., Kallergi, E., Dalezios, Y. & Tavernarakis, N. Modulation of Autophagy by BDNF Underlies Synaptic Plasticity. Cell Metab. 26, 230–242.e5 (2017).

17. Zhang, X. et al. Ischemia-induced upregulation of autophagy preludes dysfunctional lysosomal storage and associated synaptic impairments in neurons. Autophagy 17, 1519–1542 (2021).

18. Lieberman, O. J. & Sulzer, D. The Synaptic Autophagy Cycle. J. Mol. Biol. 432, 2589–2604 (2020).

19. Tomoda, T., Yang, K. & Sawa, A. Neuronal Autophagy in Synaptic Functions and Psychiatric Disorders. Biol. Psychiatry 87, 787–796 (2020).

20. Wang, Z. et al. 27-Plex Tandem Mass Tag Mass Spectrometry for Profiling Brain Proteome in Alzheimer’s Disease. Anal. Chem. 92, 7162–7170 (2020).

21. Bai, B. et al. Deep Profiling of Proteome and Phosphoproteome by Isobaric Labeling, Extensive Liquid Chromatography, and Mass Spectrometry. in Methods in Enzymology (ed. Shukla, A. K.) vol. 585 377–395 (Academic Press, 2017).

22. Ctortecka, C. et al. Comparative Proteome Signatures of Trace Samples by Multiplexed Data-Independent Acquisition. Mol. Cell. Proteomics 21, (2022).

23. Xu, S. et al. Using clusterProfiler to characterize multiomics data. Nat. Protoc. 19, 3292–3320 (2024).

24. Zhu, J. et al. Tumour immune rejection triggered by activation of α2-adrenergic receptors. Nature 618, 607–615 (2023).

25. Zhao, X., Zhang, Y., Zuo, X., Wang, S. & Dong, X. Knockdown of Adra2a Increases Secretion of Growth Factors and Wound Healing Ability in Diabetic Adipose-Derived Stem Cells. Stem Cells Int. 2022, 5704628 (2022).

26. Ali, S. & Dwivedi, Y. Early-Life Stress Influences the Transcriptional Activation of Alpha-2A Adrenergic Receptor and Associated Protein Kinase A Signaling Molecules in the Frontal Cortex of Rats. Mol. Neurobiol. (2024).

27. Sadeghi, M. A. et al. Phosphodiesterase inhibitors in psychiatric disorders. Psychopharmacology (Berl.) 240, 1201–1219 (2023).

28. Kashiwagi, E. et al. Downregulation of phosphodiesterase 4B (PDE4B) activates protein kinase A and contributes to the progression of prostate cancer. The Prostate 72, 741–751 (2012).

29. Kähler, A. K. et al. Association study of PDE4B gene variants in Scandinavian schizophrenia and bipolar disorder multicenter case-control samples. Am. J. Med. Genet. Part B Neuropsychiatr. Genet. Off. Publ. Int. Soc. Psychiatr. Genet. 153B, 86–96 (2010).

30. Chung, S. et al. Overexpressing PKIB in prostate cancer promotes its aggressiveness by linking between PKA and Akt pathways. Oncogene 28, 2849–2859 (2009).

31. Kovanich, D. et al. Sphingosine Kinase Interacting Protein is an A-Kinase Anchoring Protein Specific for Type I cAMP-Dependent Protein Kinase. ChemBioChem 11, 963–971 (2010).

32. Shaywitz, A. J. & Greenberg, M. E. CREB: A Stimulus-Induced Transcription Factor Activated by A Diverse Array of Extracellular Signals. Annu. Rev. Biochem. 68, 821–861 (1999).

33. Thiel, G. Synapsin I, synapsin II, and synaptophysin: marker proteins of synaptic vesicles. Brain Pathol. Zurich Switz. 3, 87–95 (1993).

34. Hudmon, A. & Schulman, H. Neuronal CA2+/calmodulin-dependent protein kinase II: the role of structure and autoregulation in cellular function. Annu. Rev. Biochem. 71, 473–510 (2002).

35. Koopmans, F. et al. SynGO: An Evidence-Based, Expert-Curated Knowledge Base for the Synapse. Neuron 103, 217–234.e4 (2019).

36. Subramanian, A. et al. Gene set enrichment analysis: A knowledge-based approach for interpreting genome-wide expression profiles. Proc. Natl. Acad. Sci. 102, 15545–15550 (2005).

37. Mootha, V. K. et al. PGC-1α-responsive genes involved in oxidative phosphorylation are coordinately downregulated in human diabetes. Nat. Genet. 34, 267–273 (2003).

38. Piñero, J. et al. The DisGeNET knowledge platform for disease genomics: 2019 update. Nucleic Acids Res. 48, D845–D855 (2020).

39. Langfelder, P. & Horvath, S. WGCNA: an R package for weighted correlation network analysis. BMC Bioinformatics 9, 559 (2008).

40. Pei, G., Chen, L. & Zhang, W. WGCNA Application to Proteomic and Metabolomic Data Analysis. in Methods in Enzymology vol. 585 135–158 (Elsevier, 2017).

41. Szklarczyk, D. et al. The STRING database in 2023: protein-protein association networks and functional enrichment analyses for any sequenced genome of interest. Nucleic Acids Res. 51, D638–D646 (2023).

42. Yu, Y. et al. Developmental regulation of tau phosphorylation, tau kinases, and tau phosphatases. J. Neurochem. 108, 1480–1494 (2009).

43. Li, H., Adamik, R., Pacheco-Rodriguez, G., Moss, J. & Vaughan, M. Protein kinase A-anchoring (AKAP) domains in brefeldin A-inhibited guanine nucleotide-exchange protein 2 (BIG2). Proc. Natl. Acad. Sci. 100, 1627–1632 (2003).

44. Kuroda, F., Moss, J. & Vaughan, M. Regulation of brefeldin A-inhibited guanine nucleotide-exchange protein 1 (BIG1) and BIG2 activity via PKA and protein phosphatase 1γ. Proc. Natl. Acad. Sci. 104, 3201–3206 (2007).

45. Yoon, S., Piguel, N. H. & Penzes, P. Roles and mechanisms of ankyrin-G in neuropsychiatric disorders. Exp. Mol. Med. 54, 867–877 (2022).

46. Brauchle, M. et al. Protein Complex Interactor Analysis and Differential Activity of KDM3 Subfamily Members Towards H3K9 Methylation. PLOS ONE 8, e60549 (2013).

47. Signorile, A. et al. cAMP/PKA Signaling Modulates Mitochondrial Supercomplex Organization. Int. J. Mol. Sci. 23, 9655 (2022).

48. Sunitha, B. et al. Human muscle pathology is associated with altered phosphoprotein profile of mitochondrial proteins in the skeletal muscle. J. Proteomics 211, 103556 (2020).

49. Zhong, J., Dong, J., Ruan, W. & Duan, X. Potential Theranostic Roles of SLC4 Molecules in Human Diseases. Int. J. Mol. Sci. 24, 15166 (2023).

50. Peña-Münzenmayer, G. et al. Activation of the Ae4 (Slc4a9) cation-driven Cl−/HCO3− exchanger by the cAMP-dependent protein kinase in salivary gland acinar cells. Am. J. Physiol.-Gastrointest. Liver Physiol. 321, G628–G638 (2021).

51. Kim, H. J., Kim, T., Xiao, D. & Yang, P. Protocol for the processing and downstream analysis of phosphoproteomic data with PhosR. STAR Protoc. 2, 100585 (2021).

52. Hornbeck, P. V. et al. PhosphoSitePlus, 2014: mutations, PTMs and recalibrations. Nucleic Acids Res. 43, D512–D520 (2015).

53. Kim, H. J. et al. PhosR enables processing and functional analysis of phosphoproteomic data. Cell Rep. 34, 108771 (2021).

54. Fang, X.-L. et al. Suppression of cAMP/PKA/CREB signaling ameliorates retinal injury in diabetic retinopathy. Kaohsiung J. Med. Sci. 39, 916–926 (2023).

55. Yu, L. et al. Atorvastatin inhibits neuronal apoptosis via activating cAMP/PKA/p-CREB/BDNF pathway in hypoxic-ischemic neonatal rats. FASEB J. 36, e22263 (2022).

56. Sarbassov, D. D., Guertin, D. A., Ali, S. M. & Sabatini, D. M. Phosphorylation and Regulation of Akt/PKB by the Rictor-mTOR Complex. Science 307, 1098–1101 (2005).

57. Cross, D. A. E., Alessi, D. R., Cohen, P., Andjelkovich, M. & Hemmings, B. A. Inhibition of glycogen synthase kinase-3 by insulin mediated by protein kinase B. Nature 378, 785–789 (1995).

58. Nishimura, T. & Tooze, S. A. Emerging roles of ATG proteins and membrane lipids in autophagosome formation. Cell Discov. 6, 1–18 (2020).

59. Zhao, Y. G. et al. The ER Contact Proteins VAPA/B Interact with Multiple Autophagy Proteins to Modulate Autophagosome Biogenesis. Curr. Biol. 28, 1234–1245.e4 (2018).

60. Vierra, N. C. et al. Neuronal ER-plasma membrane junctions couple excitation to Ca2+-activated PKA signaling. Nat. Commun. 14, 5231 (2023).

61. Vierra, N. C. Compartmentalized signaling in the soma: Coordination of electrical and protein kinase A signaling at neuronal ER-plasma membrane junctions. BioEssays 46, 2400126 (2024).

62. Kamemura, K. & Chihara, T. Multiple functions of the ER-resident VAP and its extracellular role in neural development and disease. J. Biochem. (Tokyo) 165, 391–400 (2019).

63. Dong, R. et al. Endosome-ER Contacts Control Actin Nucleation and Retromer Function through VAP-Dependent Regulation of PI4P. Cell 166, 408–423 (2016).

64. Yan, Z. & Rein, B. Mechanisms of synaptic transmission dysregulation in the prefrontal cortex: pathophysiological implications. Mol. Psychiatry 27, 445–465 (2022).

65. Mandal, P. K. et al. Schizophrenia, Bipolar and Major Depressive Disorders: Overview of Clinical Features, Neurotransmitter Alterations, Pharmacological Interventions, and Impact of Oxidative Stress in the Disease Process. ACS Chem. Neurosci. 13, 2784–2802 (2022).

66. Smucny, J., Dienel, S. J., Lewis, D. A. & Carter, C. S. Mechanisms underlying dorsolateral prefrontal cortex contributions to cognitive dysfunction in schizophrenia. Neuropsychopharmacology 47, 292–308 (2022).

67. Hoseth, E. Z. et al. A Study of TNF Pathway Activation in Schizophrenia and Bipolar Disorder in Plasma and Brain Tissue. Schizophr. Bull. 43, 881–890 (2017).

68. Gao, W.-J., Yang, S.-S., Mack, N. R. & Chamberlin, L. A. Aberrant maturation and connectivity of prefrontal cortex in schizophrenia—contribution of NMDA receptor development and hypofunction. Mol. Psychiatry 27, 731–743 (2022).

69. Omar, M. H. & Scott, J. D. AKAP Signaling Islands: Venues for Precision Pharmacology. Trends Pharmacol. Sci. 41, 933–946 (2020).

70. Morgan-Smith, M., Wu, Y., Zhu, X., Pringle, J. & Snider, W. D. GSK-3 signaling in developing cortical neurons is essential for radial migration and dendritic orientation. eLife 3, e02663 (2014).

71. Valvezan, A. J. & Klein, P. S. GSK-3 and Wnt Signaling in Neurogenesis and Bipolar Disorder. Front. Mol. Neurosci. 5, (2012).

72. Jope, R. S. & Roh, M.-S. Glycogen Synthase Kinase-3 (GSK3) in Psychiatric Diseases and Therapeutic Interventions. Curr. Drug Targets 7, 1421–1434 (2006).

73. Zhou, X., Wang, H., Burg, M. B. & Ferraris, J. D. Inhibitory phosphorylation of GSK-3β by AKT, PKA, and PI3K contributes to high NaCl-induced activation of the transcription factor NFAT5 (TonEBP/OREBP). Am. J. Physiol.-Ren. Physiol. 304, F908–F917 (2013).

74. Jensen, J., Brennesvik, E. O., Lai, Y.-C. & Shepherd, P. R. GSK-3β regulation in skeletal muscles by adrenaline and insulin: Evidence that PKA and PKB regulate different pools of GSK-3. Cell. Signal. 19, 204–210 (2007).

75. Peretti, D., Dahan, N., Shimoni, E., Hirschberg, K. & Lev, S. Coordinated Lipid Transfer between the Endoplasmic Reticulum and the Golgi Complex Requires the VAP Proteins and Is Essential for Golgi-mediated Transport. Mol. Biol. Cell 19, 3871–3884 (2008).

76. James, C. & Kehlenbach, R. H. The Interactome of the VAP Family of Proteins: An Overview. Cells 10, 1780 (2021).

77. Kors, S., Costello, J. L. & Schrader, M. VAP Proteins – From Organelle Tethers to Pathogenic Host Interactors and Their Role in Neuronal Disease. Front. Cell Dev. Biol. 10, (2022).

78. Bolger, A. M., Lohse, M. & Usadel, B. Trimmomatic: a flexible trimmer for Illumina sequence data. Bioinformatics 30, 2114–2120 (2014).

79. Kim, D., Paggi, J. M., Park, C., Bennett, C. & Salzberg, S. L. Graph-based genome alignment and genotyping with HISAT2 and HISAT-genotype. Nat. Biotechnol. 37, 907–915 (2019).

80. Love, M. I., Huber, W. & Anders, S. Moderated estimation of fold change and dispersion for RNA-seq data with DESeq2. Genome Biol. 15, 550 (2014).

81. Dejanovic, B. et al. Complement C1q-dependent excitatory and inhibitory synapse elimination by astrocytes and microglia in Alzheimer’s disease mouse models. Nat. Aging 2, 837–850 (2022).

82. Tan, H. et al. Refined phosphopeptide enrichment by phosphate additive and the analysis of human brain phosphoproteome. Proteomics 15, 500–507 (2015).

83. Ritchie, M. E. et al. limma powers differential expression analyses for RNA-sequencing and microarray studies. Nucleic Acids Res. 43, e47 (2015).

84. Wang, X. et al. JUMP: a tag-based database search tool for peptide identification with high sensitivity and accuracy. Mol. Cell. Proteomics MCP 13, 3663–3673 (2014).

85. Shannon, P. et al. Cytoscape: a software environment for integrated models of biomolecular interaction networks. Genome Res. 13, 2498–2504 (2003).

86. Chen, J., Bardes, E. E., Aronow, B. J. & Jegga, A. G. ToppGene Suite for gene list enrichment analysis and candidate gene prioritization. Nucleic Acids Res. 37, W305–W311 (2009).

87. Wickham, H. Ggplot2. (Springer International Publishing, Cham, 2016). doi:10.1007/978-3-319-24277-4.

88. Blighe, K., Rana, S. & Lewis, M. EnhancedVolcano: Publication-ready volcano plots with enhanced colouring and labeling. R Package Version 1, 10–18129 (2019).

89. Yang, P. et al. AdaSampling for Positive-Unlabeled and Label Noise Learning With Bioinformatics Applications. IEEE Trans. Cybern. 49, 1932–1943 (2019).

90. Dai, J., Liakath-Ali, K., Golf, S. R. & Südhof, T. C. Distinct neurexin-cerebellin complexes control AMPA- and NMDA-receptor responses in a circuit-dependent manner. eLife 11, e78649 (2022).

91. Wang, L. et al. Analyses of the autism-associated neuroligin-3 R451C mutation in human neurons reveal a gain-of-function synaptic mechanism. Mol. Psychiatry 29, 1620–1635 (2022).

